# Mapping shifts in nanopore signal to changes in protein and protein-DNA conformation

**DOI:** 10.1101/2020.04.01.020420

**Authors:** A. T. Carlsen, V. Tabard Cossa

## Abstract

Solid-state nanopores have been used extensively in biomolecular studies involving DNA and proteins. However, the interpretation of signals generated by the translocation of proteins or protein-DNA complexes remains challenging. Here, we investigate the behavior of monovalent streptavidin and the complex it forms with short biotinylated DNA over a range of nanopore sizes, salts and voltages. We describe a simple geometric model that is broadly applicable and employ it to explain observed variations in conductance blockage and dwell time with experimental conditions. The general approach developed here underscores the value of nanopore-based protein analysis and represents progress toward the interpretation of complex translocation signals.

**STATEMENT OF SIGNIFICANCE:** Nanopore sensing allows investigation of biomolecular structure in aqueous solution, including electricfield-induced changes in protein conformation. This nanopore-based study probes: (1) the tetramerdimer transition of streptavidin, observing the effects of increasing voltage with varying salt type and concentration; (2) the possible conformational states of DNA-streptavidin complexes when confined inside a pore. We describe a broadly applicable geometric approach that maps stepwise changes in the nanopore signal to real-time conformational transitions. These results represent progress toward accurate interpretation of nanopore signals generated by molecular complexes.

## INTRODUCTION

Streptavidin is a fascinating biomolecule belonging to the same family as avidin, an egg-white glycoprotein that tightly binds biotin (vitamin B7).(1, 2) The extraordinarily high affinity of streptavidin for biotin has resulted in its widespread use in a broad range of applications, including the detection of molecules in diagnostic assays,(3) the construction of nanoscale devices,(4) and the delivery of cell-specific cancer treatment.(5) In turn, researchers have investigated the properties of streptavidin and the biotin-streptavidin complex themselves, using atomic force microscopy,(6) surface plasmon resonance spectroscopy,(7) molecular dynamics,(8) and crystallography.(9, 10)

Recently, nanopore sensing and analysis have also been applied to streptavidin-based systems. For example, binding of short, biotinylated double-stranded (ds) DNA fragments with monovalent streptavidin (MSA) enabled quantification of the complex with standard nanopore electronics.(11) Further, short single-stranded (ss) DNA strands bound to MSA produced an extremely low detection rate (i.e. rate of single-molecule translocations that are detected by the bandwidth of a particular nanopore measurement system) compared to dsDNA bound to MSA,(12) allowing selective detection of RNA when bound to a complementary biotinylated strand of ssDNA complexed with MSA. Hydroxymethylcytosine was thus detected in fragmented mouse genomic DNA following addition of a biotin linker to 5-methylcytosine sites and subsequent incubation with MSA.(13) A biotinylated DNA-MSA approach was also used to identify base modifications(14) and to quantify nanomolar levels of MSA bound to 7.2 kb dsDNA.(15) More recently, MSA acted as a critical component in a scheme to detect single nucleotide polymorphism,(16) and to map the positions of short sequences in longer dsDNA.(17)

Despite the growing body of applied nanopore research into streptavidin-based complexes, no studies have compared the characteristics of passage and capture of the protein itself or the protein-DNA complex under widely varying conditions (i.e. salt type/concentration, voltage magnitude, pore size, etc.) Recent findings suggest that a finely tuned pore diameter may be critical for reproducible protein measurements(18) and that lower applied voltage *V* prevents protein elongation (|*V*| < 120 mV)(19) and time resolution artifacts such as significant broadening in the current blockage distribution (|*V*| < 200 mV).(18) Nanopore sensing of DNA-streptavidin complexes, however, has previously been performed over a range of pore sizes (7 to 9 nm) and under higher applied voltage (e.g. 600 mV),(11, 14, 15) since these conditions minimize reversible protein plugging and pore clogging. Here, we report the behavior of this versatile protein and its DNA complex under varying experimental conditions, including salt concentration (0.5 to 4 M), salt type (KCl, LiCl, NaCl) and voltage (200-600 mV). We develop a simple geometric model that is broadly applicable and discuss the proposed physical origins of various translocation signals. Our approach emphasizes the unique role of nanopore sensing in elucidating protein structure, transport dynamics and electric field interactions and offers a step forward in understanding information-rich translocation signals.

## MATERIALS AND METHODS

### Silicon nitride nanopore fabrication

In a typical experiment, a TEM grid (12 ± 2 nm thick, 0.04 mm x 0.04 mm, PN: NBPX5004Z-low res) was sandwiched between two halves of a flow cell bearing custom Viton gaskets. The flow cell was sealed shut, flushed with ethanol, then filled with 1 M KCl. The pore was fabricated using the controlled breakdown (CBD) method as described elsewhere.(20–22) Briefly, a starting bias of 5 V was applied through Ag/AgCl electrodes placed in either reservoir. The voltage was gradually increased by 0.05 V/s via custom automated software.(22) When a breakdown event was detected (through the presence of a sudden increase in current) at roughly 8 V, the software immediately removed the applied voltage and performed an IV curve between ± 200 mV. If a conductance indicative of the presence of a pore was measured, the pore was conditioned through the repeated application of 3.5 V by the software until the measured conductance *G* corresponded to 80% of the desired pore diameter. Pore diameter *d* was determined from the measured conductance value *G* using the formula 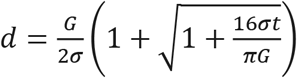, where *t* is the membrane thickness and *σ* is solution conductivity.(22) Then the voltage application was automatically reduced to 2.5 V to enlarge the pore more slowly and prevent overshooting the size. Once the target pore size of roughly 8 nm was achieved, a PSD was performed to ensure low-noise behavior. Finally, the flow cell was flushed with the solution to be used for subsequent experiments.

### Nanopore measurements and analysis

Electrolyte solution containing the analyte of interest (DNA, MSA or MSA-DNA) was introduced to the *cis* side of the custom flow cell and negative voltage was applied to the *trans* side of the flow cell in all experiments. Custom LabVIEW software was used to record translocation data using an Axopatch 200B with a 4-pole Bessel low-pass filter set at 100 kHz and a National Instruments USB-6351 DAQ card with a 500 kHz sampling rate. Origin and additional custom software were used to perform data analysis, including event level and sub-level identification through an adaptation of the Cumulative Sums (CUSUM) algorithm.(23)

Each nanopore used was first validated by recording the translocations of 1 kbp DNA. Calibration of nanopore thickness using dsDNA is described in Supplemental Information section S1, along with details of error analysis.

### Chemicals

Double-stranded DNA samples (biotinylated and non-biotinylated) were purchased from Integrated DNA Technologies (IDT, San Diego, California). The sequence of the end-biotinylated DNA fragment was as follows:

TAG A/iBiodT/T ACA GTC GTT GAC ATA GTC TCA GTT TGC TCA TTG GAA TAC ACA CCG GAT AG

Monovalent streptavidin was generously provided by the Howarth lab (Oxford, UK).(24, 25) MSA is a tetramer composed of one biotin-binding unit of hexaglutamate-tagged streptavidin (SAe) and three non-biotin-binding units of “dead” streptavidin (D). The amino acid sequences of SAe (with E6 tag underlined) and of non-biotin binding D (with residues impairing biotin binding underlined) are as follows:

#### SAe

AEAGITGTWYNQLGSTFIVTAGADGALTGTYESAVGNAESRYVLTGRYDSAPATDGSGTALGWTVAWKNNYRNAHSATTWSGQYVGGAEARINTQWLLTSGTTEANAWKSTLVGHDTFTKVK PSAAS**<u>EEEEEE</u>**

#### D

AEAGITGTWY**<u>A</u>**QLG**<u>D</u>**TFIVTAGADGALTGTYE**<u>A</u>**AVGNAESRYVLTGRYDSAPATDGSGTALG WTVAWKNNYRNAHSATTWSGQYVGGAEARINTQWLLTSGTTEANAWKSTLVGHDTFTKVK PSAAS

Streptavidin-DNA complexes were formed by incubating MSA with 56 bp biotinylated DNA in 0.3X PBS buffer for one hour at room temperature. Sample quality was confirmed through multiple electromobility shift assays (EMSA), in which biotinylated DNA was incubated with excess MSA then loaded onto a 2% agarose gel with GelRed nucleic acid stain for visualization, as described in Supplemental Information section S2.

### Images

PDB cartoons were produced using RCSB PDB Protein Workshop 4.2.0.(26)

## RESULTS AND DISCUSSION

### Monovalent streptavidin

Structural studies reveal that hydrogen bonds and salt bridges hold the four identical β-barrel subunits of a tetrameric streptavidin together.(27) Despite the symmetry of its structure, streptavidin has been called a dimer of dimers,(28) with stronger bonds existing between subunits A and B (and between C and D) (Figure 1a).(29) Across the A/C and B/D interfaces, the contact area is less than 2 nm^2^ and interactions are relatively weak.(29, 30) In wild-type streptavidin, one end of each β-barrel presents a biotin-binding pocket, meaning that a single streptavidin protein could bind up to four biotins or biotinylated molecules. Among the variety of mutant forms of streptavidin developed to limit undesired interactions, the 54.5 kDa monovalent streptavidin engineered by Howarth and coworkers(24, 25) maintains the tetrameric structure and femtomolar affinity of wild-type streptavidin for biotinylated targets, but offers a single functional biotin-binding site. Polyaspartate sequences are inserted to deactivate the biotin-binding capacity of three of the four monomers, while the addition of a hexaglutamate tag (−6e) on the final, active monomer allows for purification by ion-exchange chromatography.(25) These modifications shift the charge profile from that of wild-type streptavidin, nearly tripling its small negative charge at neutral pH.(31) The −4.2e charge of wild-type shifts to −12.2e for monovalent streptavidin, according to a basic online protein calculator.(32)

**Figure 1.**
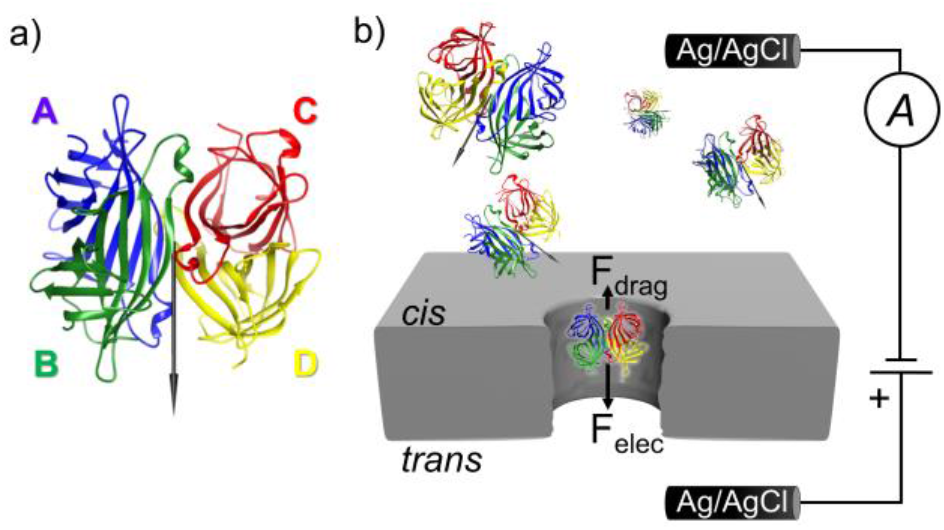
Diagrams of monovalent streptavidin and the experimental setup. a) PDB-based cartoon of monovalent streptavidin (PDB ID: 5TO2) identifying the four monomers (A, B, C and D) that make up a tetramer with stronger bonds across the A/B and C/D interfaces. The dipole moment vector (81 Debyes) is marked as a black arrow that lies in the plane of the page. b) When a positive voltage bias is applied across the nanopore membrane immersed in electrolyte solution, positive ions move through the nanopore toward the negatively charged electrode *(cis* reservoir), while negative ions and streptavidin molecules translocate through the pore toward the positively charged electrode *(trans* reservoir). The net forces acting on the molecule in the pore include contributions from a hydrodynamic drag force *(F_drag_)* generated by frictional effects such as electroosmotic flow or interaction with the pore walls and an opposing electrophoretic force *(F_elec_*).

To begin, we study the capture and translocation characteristics of MSA alone. We apply a 200 mV bias across an 8.4 nm pore in a silicon nitride membrane immersed in 4 M NaCl (pH 7.4, 1X PBS, 22.0 S/m), where the pore diameter and effective membrane thickness *H_eff_* of 11 nm have been determined using the blockage depth measured from a 1000 bp dsDNA fragment (NoLimits, Thermo Fisher Scientific, Waltham, Massachusetts),(33) as described in Supplemental Information section S1. The applied voltage drives individual streptavidin molecules from the *cis* reservoir toward the *trans* reservoir (Figure 1b). As each individual molecule passes through the pore, it blocks a corresponding volume of electrolyte solution, resulting in a measurable drop in the ionic current called an event or conductance blockage. The length of each event is referred to as its event duration or dwell time. Along with the rate of detected events, the dwell time and conductance blockage provide important information about the size, shape and charge of the translocating molecule.(19, 34–36) In this case, a 30 nM solution of monovalent streptavidin generates events (Figure 2a) at a rate of 3.13 ± 0.02 Hz (or 0.1 ± 0.01 Hz/nM) with a mean dwell time of 30 ± 20 μs and a blockage depth of 19 ± 5 nS (Figure 2b, data in red).

**Figure 2.**
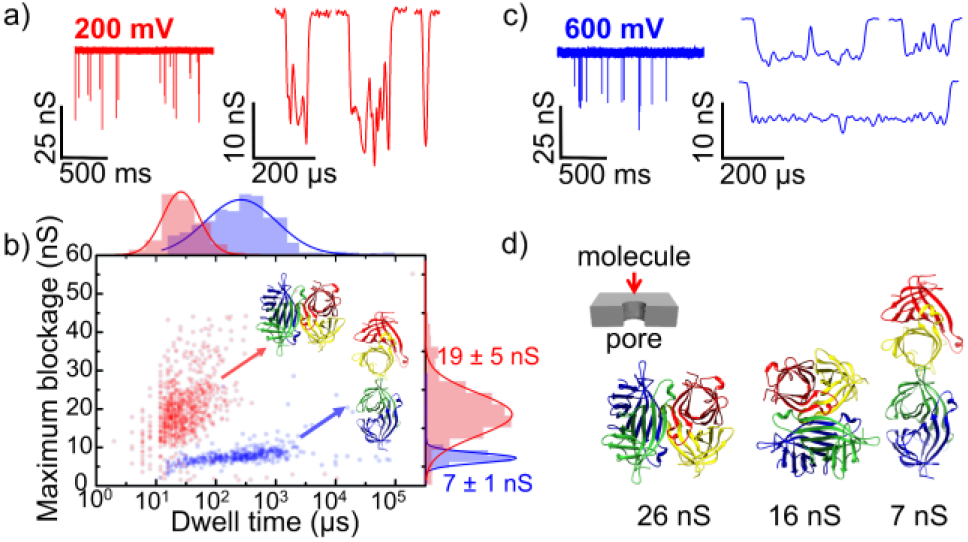
Influence of voltage on MSA signal characteristics in 4 M NaCl. a) Sample current traces for translocation of 30 nM MSA through an 8.4 nm nanopore (11-nm thick) in 4 M NaCl along with example events at 200 mV (red). b) Overlay of dwell time versus maximum blockage histograms derived from translocation signals from 900 events at 200 mV (red) and from 516 events at 600 mV (blue). At 200 mV, the mean maximum blockage depth is 19 ± 5 nS with a mean dwell time of 30 ± 20 μs. At 600 mV, the mean maximum blockage depth is 7 ± 1 nS while the mean dwell time stretches to 400 ± 400 μs. The increased dwell time at 600 mV suggests increased protein-pore interaction due to protein conformational changes, with the broad distribution reflecting the stochastic nature of translocation-induced protein deformation. Rare outlier events (which likely represent artifacts in the analysis method or brief clogging/collisions) are included to demonstrate data completeness. Insets: PDB-based cartoon of likely structures as MSA passes through a nanopore as a tetramer under an applied bias of 200 mV (red) or as two dimers at 600 mV (blue). c) Current traces and sample events for MSA translocation in the same pore at 600 mV (blue). d) PDB cartoons of MSA orientations and hypothesized structures (tetramer, split pair of dimers) corresponding to expected blockage depths calculated from the geometric model. Inset shows orientation of depicted molecules relative to pore.

In order to extract the streptavidin’s size, we must translate the measured blockage depth of roughly 19 nS into the corresponding volume of ionic solution displaced by the translocating streptavidin. Based on Ohm’s law, early work by DeBlois and Bean,(37) and the pioneering protein-sensing studies of Fologea(38) and Ledden *et al*.,(39) a folded, globular protein in its native state driven through a larger, cylindrical nanopore will generate a current change Δ*I* of:

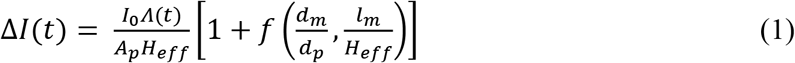

where *I*_0_ is the open-pore current, *Λ*(*t*) is the instantaneous excluded volume of the protein molecule and *A_p_* is the area of the pore. The correction factor *f* takes into account the ratio of the molecule’s diameter *d_m_* to the pore diameter *d_p_,* in addition to the molecule length *l_m_* versus the calibrated pore length *H_eff_* (which accounts for the pore’s access resistance.)(40, 41) Both the orientation and the conformation of the protein can vary during passage through the pore with significant implications for the measured current.(19, 42) For example, an unfolded protein will produce a different blockage depth from a folded protein of the same volume. Indeed, a non-spherical particle will produce a deeper blockage when its longer axis is perpendicular to the electric field in the pore.(43)

As the length of the fully folded streptavidin is well within the effective length of the nanopore *(l_m_ < H_eff_*) and the hydrodynamic diameter of the protein molecule *(d_m_* ~5.6 nm)(19, 44) is sufficiently smaller than the 8.4 nm pore diameter, DeBlois and Bean(37) show that the correction factor *f* becomes insignificant.(39) Following the work of Li,(41) Niedzwiecki(45) and coworkers, we estimate the instantaneous blockage depth for the protein to be:

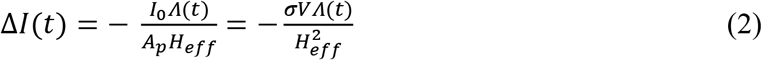

where *σ* is the measured bulk solution conductivity (22.0 S/m) and *V* is the applied potential.

Substituting our values in this particular experimental case (*V* of 0.2 V, reported volume Λ(*t*) of 94 ± 18 nm^3^ or calculated *Λ*(*t*) of 117 ± 8 nm^3^),(36, 46, 47) we obtain an expected blockage depth of 17 ± 5 nS to 21 ± 5 nS, in good agreement with the observed blockage depth of 19 ± 5 nS. Thus, we ascribe the observed blockage depth at 200 mV (Figure 2b, data in red) to the translocation of a fully folded, tetrameric monovalent streptavidin molecule.

While the volume-based model in Equation 2 accurately describes streptavidin’s translocation at low voltage, it does not address the effects of protein orientation (relative to the pore axis) or of conformational changes which become increasingly likely at higher voltages that favor protein shearing.(48, 49) Recently, Yusko(19) and Houghtaling *et al.(42)* developed a model linked to Maxwell’s derivation:

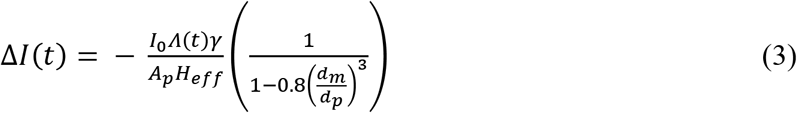

where γ is the electrical shape factor of 1.5 for a spherical molecule. In this case, the measured value of the open pore current *I*_0_ is used directly, while the area of the pore *A_p_* is calculated from the pore diameter of 8.4 nm as before. If we treat streptavidin as a sphere and substitute our values for *d_m_* of 5.6 nm, *d_p_* of 8.4 nm, and *I*_0_ of 14.2 nA, the expected blockage goes from 21 ± 5 nS to 27 ± 3 nS for the reported volume range of streptavidin. The expected blockage of 21 ± 5 nS for a spherical streptavidin molecule with a volume of 94 ± 18 nm^3^ aligns with the measured conductance of 19 ± 5 nS, suggesting again that we have observed the translocation of fully folded tetrameric streptavidin.

In reality, streptavidin is not a perfect sphere. Instead, we might consider streptavidin to be a prolate (or slightly rod-shaped) spheroid, based on the values of 4.2 nm and 5.6 nm for streptavidin’s axes reported in literature.(9, 44, 46, 50) From these values, we can determine electrical shape factors γ (as described in Supplemental Information section S3) which correspond to the extreme widthwise or lengthwise orientation of the protein relative to the pore. Substituting these values in Equation 3 produces minimum and maximum blockage depths of 16 ± 4 nS to 20 ± 5 nS for prolate length parallel to the electric field *(d_m_* = 4.2 *nm*) and 23 ± 5 nm to 28 ± 5 nS for prolate length perpendicular to the electric field (*d_m_* = 5.6 *nm*) for the range of reported volumes of streptavidin, corresponding relatively well with the measured conductance of 19 ± 5 nS, considering the associated errors.

While the models of Equations 2 and 3 succeed in estimating the conductance blockage of fully folded globular streptavidin, they become less useful in the analysis of non-spheroidal particles or multiparticle systems. Following past work,(36, 51) we seek to develop a straightforward, broadly applicable model by considering the interior of the pore as a series of two resistors: the resistance of the segment of the pore that is streptavidin-free *(R_free_*) in series with the segment of the pore containing the protein molecule (*R_with protein_*). Both the pore and the monovalent streptavidin are treated as idealized cylinders, with fully folded streptavidin occupying roughly half the length of the pore (*H_eff_* =11 nm):

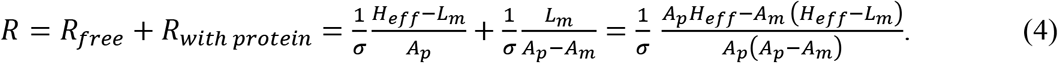

Rearranging for conductance G=R^-1^, we obtain:

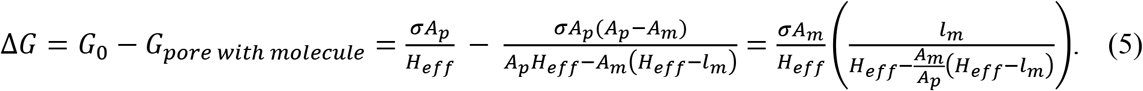

When we solve Equation 5 for this experiment (see Supplemental Information section S4), the experimentally obtained conductance blockage of 19 nS could correspond to a rod-shaped protein molecule with diameter and length of 4.8 nm (volume = 87 nm^3^), which approximates the volume of streptavidin. Interestingly, using the reported values of 4.2 nm and 5.6 nm for streptavidin’s axes, our physical model generates a more detailed picture: a blockage depth of 16 ± 3 nS for end-on translocation and a depth of 26 ±8 nS if the molecule enters the pore side-on (see Supplemental Information, section S4). Since these values represent the extremes of conductance blockage depth induced by protein orientation, we would expect the depths of the majority of events to lie between these extremes, as individual streptavidin molecules present different faces (or sample various orientations(34, 36, 42)) during their passage through the pore. The measured blockage depth amplitude and spread (19 ± 5 nS) correspond well with the values generated by our simple physical model. This correspondence between predicted and observed values for fully folded streptavidin provides preliminary experimental justification for further application of this resistors-in-series model.

Recently, Hu *et al.*(18) employed voltage-driven translocation of various dihydrofolate reductase protein mutants (L28R) to link the variation in conductance blockage amplitudes to a protein’s conformational flexibility, with a larger distribution (relative to the blockage spread obtained from a different protein) indicating a looser, more dynamically flexible shape. In the case of low-voltage streptavidin translocation under consideration here, our resistors-in-series model is a different tool, correlating the spread in blockage depth for a protein with its orientation (end-on or side-on) relative to the pore axis. The physically relevant information generated by this simple model hints at the utility of the approach, which we will apply to other translocation data obtained from MSA and MSA-DNA complexes in varying voltage and salt conditions later in this work.

To further enhance our understanding of streptavidin translocation at 200 mV, we consider dwell time *t_d_* as it relates to the protein’s charge *Q*, diffusion coefficient *D* and constant drift velocity *v.* The bulk diffusion coefficient *D* for MSA is 52.3 nm^2^/μs according to the He and Niemeyer equation(52) in 4 M NaCl (see Supplemental Information section S5). The in-pore diffusion coefficient *D(r_p_*), however, could be orders of magnitude smaller,(41, 53) even when measured using high-bandwidth equipment.(54) (Note that data collected using high-bandwidth equipment may generate a more accurate diffusion coefficient for fast translocations. When the majority of translocations cannot be detected by low-bandwidth equipment, only the longest events contribute to dwell time distributions, distorting calculated values of the protein velocity and diffusion coefficient.(55))

Haridasan *et al*.(56) recently simulated this reduced in-pore diffusion coefficient, witnessing a two-orders-of-magnitude decrease from the bulk value as streptavidin translocated through an 8 nm pore under the equivalent of roughly 500 mV applied voltage. Following the results of that molecular dynamics study, the 0.73 ratio of the MSA hydrodynamic diameter (6.2 nm, see Supplemental Information S6 for calculation) to the pore diameter (8.4 nm) may cause the protein translation time to deviate from the frictional drag relationship ~*r_h_*/(*r_p_ – r_h_*) expected in larger pores.(57) Instead, interactions between MSA and the nanopore interior will produce excessive drag, reducing the streptavidin’s mobility to a value that depends on the applied force.(56) Pore-protein interactions likely produce the dramatically prolonged dwell times (30 ± 20 μs) observed here.

Protein mobility *μ* in nanopore translocation experiments is routinely described(38, 43, 54) with the Einstein-Smoluchowski equation:

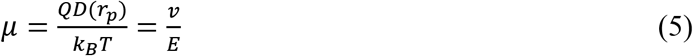

where *Q* is the protein charge, *D*(r_p_) is the diffusion coefficient within the pore, *v* is the protein velocity, *E* is the electric field, and *k_B_T* is the product of the Boltzmann constant and temperature (a scale factor for energy values equivalent to roughly 4 pN·nm). Using the common assumption(38, 43, 54) that voltage drops entirely across the effective length of the pore (*E* = |*V*|/*H_eff_*), the measured mean dwell time of 30 ± 20 μs corresponds to an average constant drift velocity *v* (calculated as *H_eff_/t_d_*) of 0.4 ± 0.2 nm/μs through this 8.4 nm diameter pore under our experimental conditions. Rearranging the equation and substituting the charge of −12.2e for monovalent streptavidin, we obtain a value for the in-pore diffusion coefficient *D*(*r_p_*) of 4.3 x 10^-2^ nm^2^/μs, which is roughly 3 orders of magnitude less than streptavidin’s bulk diffusion coefficient *D* of 52.3 nm^2^/μs in 4 M NaCl. An in-pore translational mobility *μ* of 20 nm^2^μs^-1^V^-1^ for streptavidin is also easily found from (5). The significant reduction in the values for in-pore diffusion coefficient and corresponding mobility from the bulk values is in line with prior studies(41, 53, 54, 56) and largely attributable to the effects of confinement and strong interactions between the streptavidin and the pore walls.

Interestingly, Muthukumar(53) has offered an adaption of Equation 5 which takes into account nanopore-related discrepancies such as the counterion cloud, confinement effects and protein-pore interaction. This generalization can be described as follows:

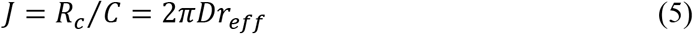

where *Q*(*r_p_*) and *D*(*r_p_*) are the effective in-pore values for the charge and the diffusion coefficient for the protein. Of interest is the effective charge *Q*(*r_p_*), equivalent to *Q*/(1 + *κr_g_*), which accounts for the drag force on the protein from the counterion cloud within the pore. Using the salt-concentration-dependent inverse Debye length κ and the protein’s radius of gyration *r_g_*, we find an effective charge *Q*(*r_p_*) of 0.79e, a value that is ~7% of the MSA charge of −12.2e (see Supplemental Information section S7 for detailed calculation). Assuming once again that the applied voltage drops across the effective length of a pore and that protein velocity *v* exhibits a linear dependence on voltage, an in-pore diffusion coefficient *D*(*r_p_*) of 0.66 nm^2^/μs is produced by rearranging Equation 5 to *D*(*r*) = *vH_eff_k_B_T*/*VQ*(*r_p_*) where *v* = 3.7×10^-4^ m/s, *Q*(*r_p_*) = 0.79e, *k_B_T* = 0.0257 eV, *H_eff_* = 11 nm. At roughly 1% of the bulk constant, this value is in good agreement with the value predicted by the Bowen, Mohammad and Hilal expression(58, 59) valid for a ratio of the hydrodynamic radius of the protein to the pore radius (*r_h_/r_p_*) up to 0.8 (see Supplemental Information section S8 for calculation), which generates an in-pore diffusion coefficient that is <2% of the bulk value for a particle in a cylindrical pore.

Although pore-related effects can explain the reduced in-pore diffusion coefficient, missed events may also skew our results. Missing a significant number of fast translocations would reduce the measured average velocity and the associated diffusion coefficient. The Axopatch 200B patch-clamp amplifier used here may detect only those events delayed by protein–pore interactions or lying in the tail of the residence-time distribution,(55) with a significant portion of the streptavidin molecules translocating too quickly for detection (see Methods section.) Evidence of some missed events can be seen in the sharp cutoff as dwell times approach 10 μs in Figure 2b (data in red), where each data point represents the conductance blockage and dwell time of a single translocation.

One approach to confirm missed events(54, 55, 60, 61) employs the Smoluchowski equation which describes the rate of protein arrival *J* at a perfectly absorbing hemisphere of radius *r_eff_*:

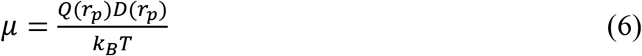

where *R_c_* is the detected capture rate, *C* is the protein concentration, *D* is the bulk diffusion coefficient.(62–65) The voltage-dependent hemisphere radius *r_eff_* is typically estimated to be the radial distance away from the pore where the electrophoretic and diffusional forces are equal. Rearranging the equation to obtain r_eff_ = R_C_/(2πDC) and substituting our values at 200 mV (R_C_ = 3.13 s^-1^, C = 30 nM or 1.8 x10 molecules/m, D = 52.3 x10^-12^ m^2^/s), we arrive at an effective radius of 0.5 nm, or roughly 12% the size of the actual pore radius of 4.2 nm. Using similar equipment and a larger nanopore (10 nm radius), Plesa *et al.* saw an even smaller capture radius for streptavidin: roughly three orders of magnitude smaller than the pore radius.(55) However, high-bandwidth equipment and a smaller pore size (2.6 nm radius) with an applied voltage |*V*| < 125 mV permitted Larkin *et al*.(54) to demonstrate that capture radii can extend up to ~ 20 times beyond the pore radius when fast protein translocations are not missed by the detector.

We might expect a capture radius larger than the pore radius due to the enhanced capture of charged biomolecules in an applied electric field, but the complete picture is more complex. The biomolecule may experience strong currents from electroosmotic flow, in addition to the electrophoretic force associated with its charge. These electrophoretic and electroosmotic forces have opposing effects on capture rate for negatively charged streptavidin molecules approaching the negatively charged SiN pore,(61) which may present additional barriers such as loss of conformational entropy in its restrictive space and interactions with its walls.(66) Even for proteins with a greater negative charge than that of monovalent streptavidin, the combination of these effects can shrink the capture radius below the pore radius in the barrier-limited regime.(61) The fact that our calculated capture radius and the pore radius are of the same order of magnitude suggests that we detect a significant portion of the total translocation events. Even if our bandwidth-limited equipment principally detects events delayed by protein–pore interactions,(55) sufficient relevant data is collected to observe trends, to draw comparisons across various experimental conditions and to calculate meaningful values, such as the in-pore diffusion coefficient.

At this point, we have thoroughly investigated the transport characteristics of streptavidin at 200 mV through an 8.4 nm diameter nanopore of 11 nm thickness and have demonstrated that the detected events correspond to translocation of the tetrameric macromolecule. Nevertheless, a simple increase in the applied voltage to 600 mV across the same nanopore yields surprising results in the detected events (Figure 2c). The mean blockage depth abruptly drops to 7 ± 1 nS (Figure 2b, data in blue). This shift in measured conductance cannot be attributed to mere signal attenuation due to bandwidth limitations since the average dwell time of detected events simultaneously increases to roughly 400 μs, well above the temporal resolution of the sensing equipment (Figure 2b, data in blue). Tripling the voltage also triples the capture rate to 9 ± 1 Hz (or 0.3 Hz/nM).

The volume-based blockage depth Equation 2 suggests that the protein volume has decreased to around 39 nm^3^, or 30 to 40% of the original volume (94 to 117 nm^3^). Since the dwell time of detected events is roughly 10 times more than the dwell time at 200 mV, it seems unlikely that the protein is simply compressed significantly in both dimensions, as that would produce a shorter event. Prior publications show decreasing blockage depths for increasing voltage due to electric-field-induced stretching or unfolding of proteins.(67, 68) Although experimental studies(41, 43, 69–72) and simulations(73, 74) have reported that electric field alone can compromise protein structural integrity, rupturing MSA requires an opposing force. As the electric field drives the protein through the pore, strong interactions between the molecule and the pore walls likely cause protein rupture, resulting in a dramatic drop in excluded volume (i.e. maximum blockage) and a significant increase in mean dwell time (Figure 2b, compare data in red and blue). The stochastic nature of translocation-induced protein deformation produces a broader distribution of dwell times,(49) with anomalously long events corresponding to newly exposed residues interacting with the pore.(43) Thus, the most likely explanation for the decreased blockage depth and increased dwell time is a partial unfolding or restructuring of the protein under the larger force of a 600 mV applied bias.

Indeed, the structure of streptavidin can be disrupted thermally,(75) or by mechanical stress.(29) An atomic force microscope (AFM) applying a stretching force of 100 pN at the pulling speed of 500 nm/s consistently ruptures the 4 hydrogen bonds binding the two dimers of streptavidin together.(76) Similarly, a recent theoretical study modeled the effects of a nanopore’s electric field on streptavidin and found potential changes in streptavidin’s secondary structure under certain pulling conditions at 2 V through a 10-nm diameter hour-glass-shaped nanopore in 1 M KCl.(77)

The sequence of residues composing monovalent streptavidin contribute a total charge of −12.2e at neutral pH, but the net charge does not give the full picture. The non-uniform nature of the charge distribution in proteins renders them highly susceptible to electrophoretic effects: negatively charged portions of the protein are pulled in the opposite direction from positive portions, resulting in stretching(67) or rupture in the case of sufficiently high voltages.(43, 71, 78) The heterogeneity of the charge may also lead to slowing of translocation at specific points.(41) Further, electro-osmotic effects, driven by a flow of counterions along the charged pore wall, can counteract or even reverse electrophoretic flow.(79)

MSA will likely favor entry into the pore at the negatively charged hexaglutamate tag on its C-terminal. Once in the pore, the protein will sample a variety of orientations relative to the electric field, which will exert a torque on the protein’s dipole moment. The alignment of a protein’s dipole moment is a superposition of the individual dipoles of each of its N amino acids.(70, 80) Although each monomer of wild-type streptavidin has a calculated dipole moment of 256 D according to the Weizmann server,(81, 82) the individual moments of the monomers cancel each other out when aligned in the tetramer.(19, 42, 83)

Such cancellation of the individual moments of the monomers does not occur for monovalent streptavidin. Slight variations in the structure of MSA compared to that of wild-type streptavidin introduce a dipole of around 80 D,(81, 82) as seen in Figure 1a. Thus, the MSA will experience torque-induced rotation and a net force in the non-uniform electric field near the nanopore. Following the work of Talaga *et al.*(41) and Freedman *et al*.,(43) forces acting on opposite charges within the protein may displace its chains from their native positions and further enhance the molecule’s effective dipole moment, increasing its magnitude in higher electric fields. Monovalent streptavidin contains 45 negative charges (6 of them localized to the hexaglutamate tag on its C-terminal) and 32 positive charges that are driven in opposite directions by an applied voltage. The force per unit charge at 600 mV would be ~6 pN, suggesting sufficient force available in the system to approach the streptavidin-rupturing values observed by AFM.(76) In comparison, electrophoretic forces reaching ~100 pN on DNA in nanopores have been reported for applied voltages > 300 mV.(84–89) Mapping the force distribution on the protein is beyond the scope of this study, as it would require consideration of numerous factors, including charge screening effects from the ionic solution,(90, 91) interactions with the pore walls, electroosmotic flow,(79, 92) and the possibility of a significantly smaller unfolding force compared to that of AFM.(93)

The 10 mV/nm field in our experiment likely disrupts the antiparallel beta barrel tertiary structure of the MSA by breaking the hydrogen bonds along the weakest plane of the MSA dimers. MD simulations suggest that a static external field with twice the strength is sufficient to radically alter protein conformation, destabilizing α-helical structures and triggering the formation of β-sheet secondary structures.(94) Similar structural rearrangement due to enhanced polarization of protein molecules in a 10^8^ V/m field has also been observed.(95)

We therefore propose that the decrease in blockage depth and concomitant increase in dwell time at 600 mV result from rending the streptavidin tetramer in half then pulling the pieces through the pore in rapid succession (Figure 2d). A significant mid-event current drop (to less than half the maximum blockage amplitude) is visible in roughly 30% of all events (and in over 60% of events longer than 1 ms), likely corresponding to the brief interval measurable by our electronics before the second half of the streptavidin tetramer follows the first. Interestingly, when we apply the resistors-in-series calculation, we obtain the observed conductance blockage of 7 ± 1 nS from a cylinder with diameter 2.1 nm (half the 4.2-nm width of the dimer on one axis) that occupies the entire pore length (see Supplemental Information, section S4). This could be the case if the two dimers do not completely detach or if the second dimer rapidly follows the first. If the streptavidin splits on the longer axis (5.6 nm), we obtain blockage depths of 5 ± 1 nS and 9 ± 2 nS for a 4.2 x 2.8 nm dimer, depending on its orientation. Again, the simple geometric model makes physical sense and helps us form a hypothesis for the physical origins of the measured signal. While typical models either ignore particle orientation (volume-based model of Eq. 2) or assume spheroidal particles (electrical shape factor model of Eq. 3), the resistors-in-series approach described here expands the existing toolkit by enabling analysis of non-spheroidal particles (e.g. partially folded proteins) and multi-component systems (e.g. DNA-protein complexes).

We have elucidated the behavior of streptavidin in 4 M NaCl at low and high voltages in a single pore (8.4 nm diameter, 11 nm thick), observing the crossover from folded tetramer to dimer translocation configuration. We next explore a few combinations of cationic species and ionic strengths, comparing the streptavidin translocation characteristics in 2 M KCl, 2.9 M LiCl and 4 M NaCl. (Detecting sufficient streptavidin translocations at the same concentration in different salts proved challenging due to a combination of factors, including sub-resolution dwell times, low capture rate, pore clogging and unintended variation in pore size.) All these concentrations lie in the high-salt regime, where the Debye length is small compared to the pore radius and charge screening should be similar. The effect of each ion type depends on its specific interactions with the protein backbone and charged side chains in solution.(96–98) For example, high NaCl concentration destabilizes both spider silk dimers(99) and beta clamp dimers,(100) but confers enhanced stability to formyltransferase dimers.(101) Whether increasing or decreasing protein stability, however, these chloride salts act in a roughly linear fashion with increasing concentration.(102)

We first collect data with a similarly sized pore (9.6 nm diameter, 11 nm thickness) in 2 M KCl (pH 8, 5 mM HEPES, 0.03X PBS, 18.4 S/m). This time, applying 200 mV across the pore produces no detectable translocations (see Supplemental Information section S9). However, at 800 mV, events become visible at a rate of 0.4 Hz/nM with blockage depths of roughly 4 ± 1 nS and dwell times centered around 30 ± 20 μs. This blockage depth corresponds to a volume of roughly 36 nm^3^, ~1/3 of the full streptavidin volume, in close agreement with the value observed for streptavidin dimers in 4 M NaCl at 600 mV. It therefore appears that the streptavidin is separated into dimers at 800 mV in 2 M KCl. We again attempt to observe translocations at 200 mV using two similarly sized pore (8.6 nm diameter x 18 nm thick, 9.4 nm diameter x 11.7 nm thick) but detect no events, suggesting that the streptavidin molecule passes through the pore too rapidly at 200 mV in 2M KCl. (Hindered passage is unlikely here since both pores are larger than the 8.4 nm diameter pore through which clear events were detected in 4 M NaCl, as described above.)

Despite detecting no events in a series of pores at 200 mV in a 2 M KCl solution containing streptavidin, we adjust the salt concentration to 0.5 M KCl (5.9 S/m) and detect events in an 8.4-nm pore (8.7-nm thick). The measured rate of 300 ± 20 Hz for a 1200 nM MSA solution (or 0.25 Hz/nM) with the 200 mV applied bias corresponds to an effective capture radius of roughly 0.8 nm (see Supplemental Information S10 for calculation); as discussed earlier, the small size of this capture radius compared to the 4.2 nm pore radius suggests that our bandwidth limited equipment may miss fast translocation events and/or that we are operating in a barrier-limited regime. The average dwell time is roughly 100 μs with a broad maximum blockage depth peak from roughly 2 to 4 nS, with a mean value of 3 ± 1 nS (Figure 3a, data in brown). The volume model generates a value of 25-50 nm^3^ for these blockage depths, or roughly 30 to 50% of the full streptavidin volume. As depicted in Figure 3b, the geometric model reveals that a streptavidin dimer measuring 2.8 x 4.2 produces a 2 nS blockage, while two dimers extended over the length of the pore (i.e. translocating in rapid succession) would produce a depth of 4 nS. Thus, we see that the streptavidin, in the confined environment of the pore, is reduced to dimer form at 200 mV in 0.5 M KCl, whereas it remained fully folded at 200 mV in 4 M NaCl.

**Figure 3.**
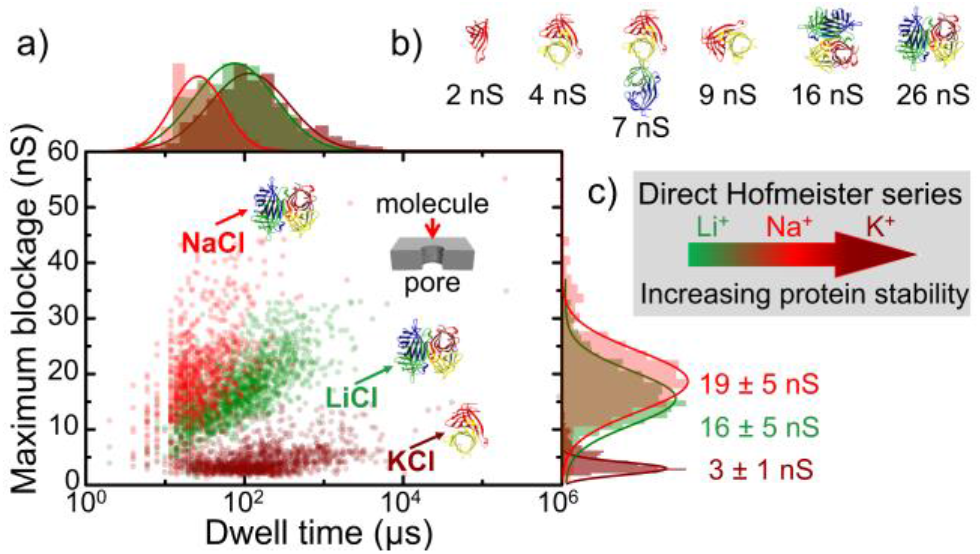
Nanopore detection of monovalent streptavidin (MSA) in various salts at 200 mV. a) Overlay of maximum blockage and dwell time histograms derived from translocation of MSA. Green dots correspond to individual events from translocation of MSA in 2.9M LiCl (pore: 7.4 nm diameter x 8.7 nm thickness), while red dots correspond to 4 M NaCl (pore: 8.4 nm diameter x 11 nm thickness) and brown dots are from translocations in 0.5 M KCl (pore: 8.4 nm diameter x 8.7 nm thickness). Inset shows orientation of depicted molecules relative to pore. b) PDB cartoons of MSA orientations and hypothesized structures (tetramer, dimer, monomer) corresponding to expected blockage depths calculated from the geometric model. c) Depiction of cationic Hofmeister series as arrow provides qualitative ranking of ions with respect to their power to increase protein stability with Li^+^ < Na^+^ < K^+^.

Finally, we test the effects of Li^+^ on streptavidin translocation by applying a 200 mV bias across a 7.7 nm pore (8.7 nm thickness) in 2.9 M LiCl (pH 8, 0.2X PBS, 14.9 S/m). For a concentration of 98 nM, the event rate is 6.4 ± 0.4 Hz (0.06 Hz/nM), suggesting a capture radius of roughly 0.3 nm. This relatively small capture radius indicates some undetected fast events, a barrier-limited regime, or a combination of both factors. The mean dwell time is around 70 μs with conductance blockage peaking at 16 ± 5 nS (Figure 3a, green), which corresponds to a volume of 86 nm^3^, matching the volume obtained in 4 M NaCl at 200 mV. The geometric model suggests that a tetrameric streptavidin (4.2 x 5.6 nm) entering the pore on its shorter axis corresponds to 16 nS (Figure 3b). The range of observed blockages could correspond to a number of physical states: two dimers extended through the pore delivers 10 nS of blockage depth, as does a single dimer traveling through the pore sideways (5.6 nm axis perpendicular to the pore walls). In any case, the vast majority of events correspond to the fully folded form of streptavidin.

It is intriguing that a 200 mV bias appears sufficient to separate the streptavidin dimers in 0.5 M KCl, but not in 2.9 M LiCl or 4 M NaCl. Although an adequate number of events could not be gathered in each salt at the same concentration due to experimental challenges (e.g. sub-resolution dwell times, low capture rate, pore clogging, unintended variation in pore size), we can consider potential effects of the combined changes in salt concentration and type. Research into protein-specific salt effects has intensified(96, 98, 103, 104) since the Hofmeister (or lyotropic) series first ranked salts according to protein-stabilizing efficiency (Figure 3c).(105, 106) Nevertheless, the behavior of specific proteins in salt solutions remains complex and difficult to predict.(96, 107, 108) We will examine three potential mechanisms of interaction between the streptavidin protein and 1) the anionic portion of the salt, namely Cl^-^, 2) the cationic portion of the salt (K, Na or Li) and 3) the salt concentration (0.5, 2.9 or 4 M).

At first glance, we might expect the most significant Hofmeister effects from 1) the anionic portion of the salt, since anions tend to interact more closely with the protein than do cations which cannot easily access the highly solvated anionic groups typical of biomacromolecules (e.g. glutamate).(107) In this case, the 4 M NaCl solution with its large concentration of mildly chaotropic Cl^-^ anions could exert the strongest influence by interacting with hydrogen-bond donor groups and destabilizing the protein’s folding.(109) However, we observe the opposite trend in Figure 3a: the protein is most stable in the highest concentration of Cl^-^ ions (4 M NaCl) and least stable (i.e. undergoes partial unfolding during translocation) in the lowest concentration of Cl^-^ ions (0.5 M KCl). To explain this deviation, we look to a second potential effect: the weaker interaction of 2) the cationic portion of the electrolyte with anionic portions of the streptavidin. The strongest effect on the protein here could arise from the small but highly hydrated lithium cations, which tend to destabilize folding for negatively charged proteins like monovalent streptavidin by interacting favorably with amide groups (i.e. the protein backbone.)(96) Again, the opposite trend appears in Figure 3a: the brief deep events associated with the LiCl solution suggest the protein remains folded, while the long shallow events in KCl suggest partial unfolding, despite the prediction that K^+^ cations could promote structural stability for negatively charged proteins.(96) The explanation for the observed behavior may lie in the final effect we will consider: 3) salt concentration. Whether a specific salt enhances or diminishes a protein’s structural stability, its effects are roughly linear with increasing concentration in the moderate-to high-salt regime.(96, 102) However, at very high concentrations (> 4 M), salts can enhance protein stability (while decreasing its solubility) through volume exclusion.(96) The mechanism is not fully understood,(110, 111) but it appears that excessive salt ions in solution interact preferentially with water, leaving fewer water molecules available for the protein. The hydrophobic forces that drive protein folding are strengthened, potentially “squeezing” the protein to the point of aggregation and precipitation.(96, 107) If higher salt concentrations produce more compact, stable protein folding, we have a potential explanation for faster events corresponding to increasing salt concentration. The order of decreasing event duration (KCl > LiCl > NaCl) corresponds to increasing ionic strength from 0.5 M KCl to 2.9 M LiCl to 4 M NaCl. The longest events occur in 0.5 M KCl, where the relative drop in measured blockage amplitude confirms partial unfolding of the protein, while much shorter, deeper events at higher salt concentration correspond to translocation of the fully folded protein. The shift from 0.5 M KCl to higher salt concentrations (2.9 M LiCl or 4 M LiCl) appears to promote protein folding, reinforcing the streptavidin tetramer enough to prevent its dissociation at 200 mV. It is worth noting that while this volume-exclusion argument could explain the observed phenomena, it neglects other potential contributions, such as the effects of the Hofmeister series on electroosmotic flow(112) and on protein-pore interaction.(113) We leave a deeper consideration of streptavidin-salt interactions to future studies, which will include data collected in different salts at the same concentration.

To summarize the results of our nanopore sensing of MSA, we observe translocation of the fully folded protein in 4 M NaCl at 200 mV, while the same protein seems to undergo a tetramer-dimer transition at 600 mV. The event rate shows a linear dependence on voltage, but a portion of the events may not be resolved by our bandwidth-limited equipment. In 2 M KCl, no protein translocation is visible at lower voltages, but dimer pairs appear again at 800 mV. At a KCl molarity of 0.5 M, we observe streptavidin dimer translocations even at a relatively low voltage of 200 mV, possibly because the destabilizing effect of the electric field in the pore is leveraged by the protein’s reduced folding stability at this lower salt concentration. At the same voltage, increasing ionic strength allows observation of fully folded streptavidin in 2.9 M LiCl as in 4 M NaCl. The lowest event rate of 0.06 Hz/nM at 200 mV occurs in 2.9 M LiCl, while 2 M KCl offers the highest of 0.4 Hz/nM at 800 mV.

### MSA bound to biotinylated dsDNA

Equipped with a better understanding of the physical origins of electrical signatures observed during the translocation of streptavidin alone, we turn our attention to the characterization of monovalent streptavidin-DNA complexes. For this, we first incubate MSA with end-biotinylated 56 bp doublestranded DNA. A shift in the total volume of unoccupied streptavidin from 105 ± 3 nm^3^ to 133 ± 2 nm^3^ upon biotinylation was measured by Neish *et al.* using AFM.(6) Although biotin is a small molecule, it induces a 1 to 1.5 nm shift in streptavidin when the loop in streptavidin’s binding pocket closes over it, in addition to a very slight contracture in the overall streptavidin structure.(114) Neish proposed that this contracture might “harden” the protein, reducing its deformability by the AFM tip and producing a larger volume measurement.(6) To account for this slight increase in volume (or hardness) upon biotinylation, we select larger values from the range reported for biotin-bound streptavidin, 5.8 x 4.5 nm,(44, 47, 115) compared to the dimensions of 5.6 x 4.2 nm used for the streptavidin alone.

We return to the same nanopore previously used to measure 30 nM MSA alone in 4 M NaCl (Figure 2) and introduce a 4 M NaCl solution containing 30 nM end-biotinylated 56 bp dsDNA pre-incubated with 30 nM MSA into the *cis* reservoir (see Methods section). (As observed in prior nanopore studies,(11, 12) translocation of the 56 bp dsDNA alone produces no events detectable by the Axopatch 200B since short DNA fragments traverse the pore so quickly that smaller pores (<4 nm) and faster electronics are needed to fully resolve these fast events.(116)) With 200 mV of applied voltage, we observe MSA-DNA events such as those in Figure 4a. The measured rate of detected events is nearly double that of bare MSA under the same conditions, while the dwell time increases from 30 ± 20 μs for bare MSA to 900 ± 300 μs after binding to biotinylated DNA. At the same time, the single maximum conductance blockage depth of 19 ± 5 nS for bare MSA becomes two deeper peaks for the MSA/DNA complex, with a sharp peak at 52 ± 1 nS and a broader secondary peak at 45 ± 5 nS (Figure 4b). Since we use the same pore that maintains a stable conductance for the entire experimental set, these shifts cannot be attributed to pore-to-pore variability (e.g. pore thickness, shape or surface charge.)

**Figure 4.**
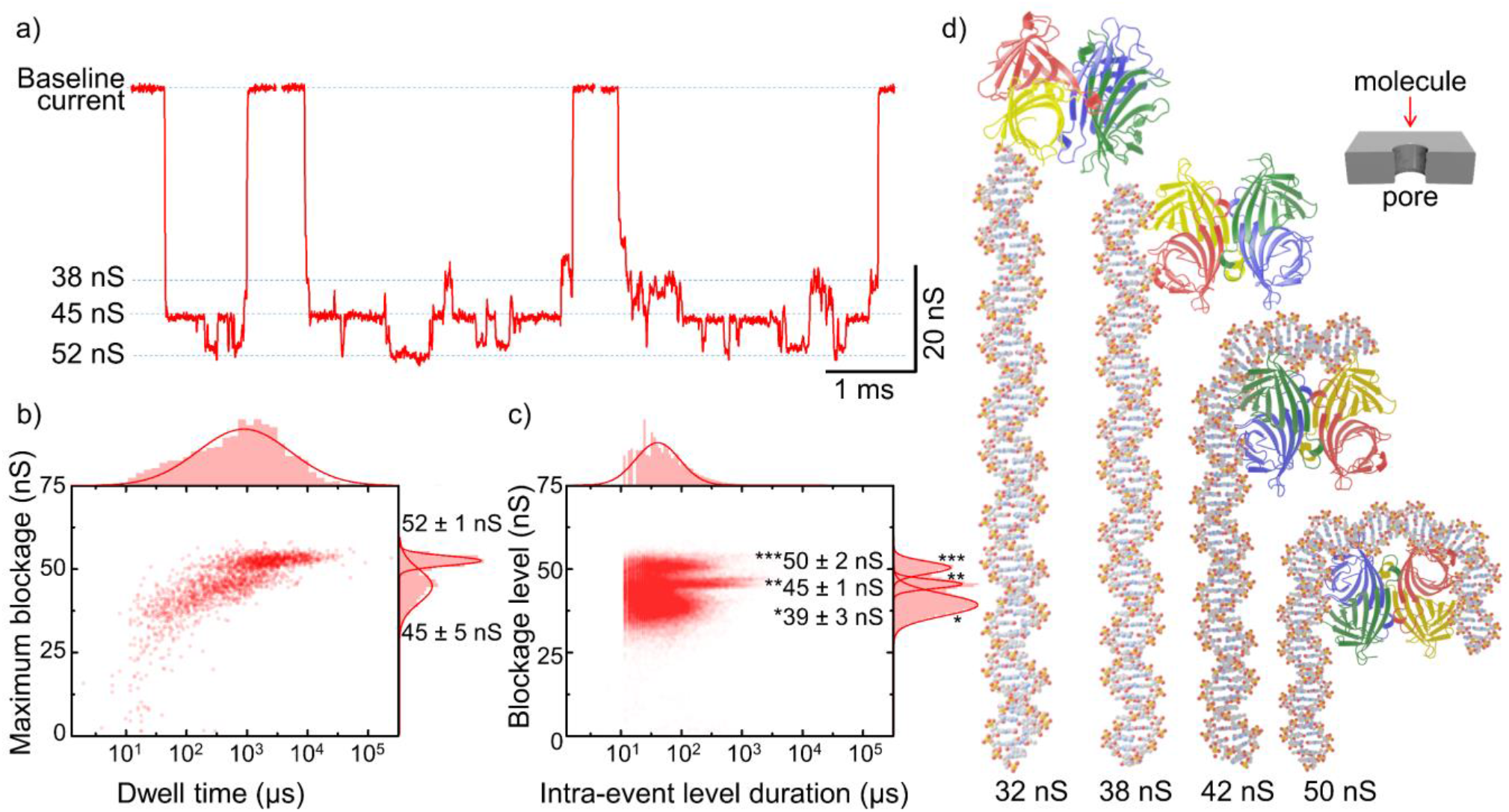
Nanopore detection and analysis of MSA bound to biotinylated dsDNA in 4 M NaCl at 200 mV. Data obtained using the same pore as in Figure 1 (8.6 nm diameter, 11 nm thickness). a) Sample event traces. b) Overlay of dwell time and maximum blockage histograms. c) Overlay of intraevent level duration versus blockage levels for the same data. d) PDB-based cartoons of hypothesized MSA-DNA orientations. Inset shows orientation of depicted molecules relative to pore.

Closer examination of the current trace reveals discrete transitions in the current level within individual events (Figure 4a). In fact, an average of thirty transitions are detected in each of the ~1900 events. Although about half of the events achieve the deepest level (52 nS), the deepest level does not occur on its own: all events containing the deepest level also contain significantly lower levels (e.g. 37 nS). An overlay of intra-event blockage level versus level duration histograms shows three principal intra-event blockage depths at roughly 40, 45 and 50 nS (Figure 4c), with the mean minimum blockage value for all events at 34 ± 3 nS. (The mean minimum blockage is extracted from a single peak Gaussian fit to the minimum blockage depth histogram with the error representing the standard deviation extracted from the same fit.) Only about 10% of the total events display a single blockage level, with the value of that level measuring 37 ± 4 nS. Rather than being the exception, long events with discrete levels are the rule, and similar discrete intra-event levels are also detected in LiCl and KCl for pores of similar diameter (see Supplemental information section S11).

Parenthetically, large intra-event amplitude fluctuations of ~7 nS in the blockade current have previously been reported by Shi *et al*.(117) when streptavidin was used as a molecular plug in a 3 to 5-nm diameter pore with 10 nm thickness at 100 mV. In that case, the streptavidin was too large to translocate through the pore. The authors attributed the long and sporadic motion of the docked DNA-streptavidin to non-specific adhesion with the nanopore surface, which allowed the complex to adjust and reorient relative to the pore. While we see blockage depths that fluctuate between discrete levels for streptavidin-DNA complexes, our case differs in three important ways: (1) our 8.6-nm pore is more than large enough to allow passage of fully folded monovalent streptavidin at 200 mV as confirmed in earlier experiments, (2) our observed blockage depths for MSA-DNA are significantly deeper than those observed by Shi *et al.,* corresponding to full passage rather than temporary docking of the complex and (3) our events self-resolve and thus do not require flipping the polarity of the applied voltage to release the complex.

To shed light on the physical meaning of the detected blockage depths, we plug the mean blockage depth of single-level events (37 nS) into the simplest volume Equation 2 and generate a value (200 ± 40 nm^3^ ) corresponding to the combined volumes of the biotin-bound tetrameric streptavidin (133 ± 2 nm^3^)(6) and the entire length of the DNA fragment (72 ± 8 nm^3^). However, it is not apparent from this model how the entire 19-nm length of DNA would fit into the 11-nm length of the pore along with the streptavidin. Further, the larger blockage depths of 45 and 52 nS have no physical meaning, since they correspond to volumes larger than the maximum combined volume of DNA and streptavidin.

Employing the electrical shape factor approach of Equation 3 offers a clue. For a spherical particle (γ = 1.5) with the maximum combined volume of DNA and MSA, we obtain a blockage depth of 34 ± 6 nS when the correction factor is set to 1 (as when the nanopore diameter is much larger than the longest dimension of the protein). However, the correction factor must be considered since the longest axis of the protein (5.8 nm) approaches the diameter of the pore (8.6 nm). With the correction factor, a spherical particle of diameter 6.5 nm with the combined volume of MSA-DNA produces a 52 ± 7 nS drop. For this deeper blockage, the electrical shape factor model suggests a spheroidal particle with the effective combined diameter and volume of MSA plus DNA, implying that the DNA is bending around the streptavidin (see Supplemental Information section S12, case 3 for calculation.)

Bending of the short dsDNA fragment may defy expectation because bare dsDNA shorter than its persistence length (reportedly ~40 to 50 nm)(118–120) is often treated as a rigid rod in aqueous solution.(121) Nevertheless, DNA can bend spontaneously(122) and behave with surprisingly high flexibility even in short segments.(123) Following Garcia *et al*.,(124, 125) the force required to bend a 56 bp fragment of dsDNA into an arc of circle with radius 2.9 nm (corresponding to the long axis of biotin-bound streptavidin) can be described as:

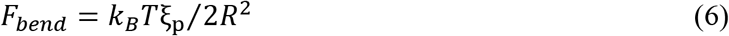

where ξ_p_ is the persistence length (taken as 40 nm for DNA)(118) and *R* is the bending radius. Substituting these values reveals that a ~10 pN force is sufficient to bend the dsDNA tether around the MSA, though less force may be required in high salt conditions.(126, 127) With an applied voltage of 200 mV in this 11-nm thick nanopore, we find from the relation *E* = |*V*|/*H_eff_* that the electric field strength is roughly 1.8 x10^7^ V/m with a corresponding force per unit charge of ~2 pN. MSA bears a charge of −12.2e, with 6 charges localized to the hexaglutamate tag on the active subunit of the protein (along with the tetramer’s 81 D dipole moment). Even a 60% reduction in the effective MSA charge due to screening in ionic solution(90, 91, 128) would leave sufficient force available in this system to bend the DNA around the protein, although a detailed picture of the forces acting on the DNA-protein complex is beyond the scope of this study.

As for the discrete intra-event blockage levels, the resistors-in-series model provides further clarification. Translocation of the DNA-protein complex generally begins with a blockage depth between 30 and 38 nS (Figure 4a). This range suggests that the DNA fragment is entering the pore with the width-wise streptavidin trailing behind or just alongside the tail end of the fragment (Figure 4d). Judging from the long-lived events observed in the current trace, the DNA-streptavidin complex experiences significant interactions with the interior surface of the pore, providing time for both the MSA and the dsDNA tether to adjust and reorient. This non-specific adhesion to the nanopore interior may stall the protein-DNA complex, while the driving electrophoretic force induces conformational changes. Applying the resistors-in-series model to the observed discrete intra-event blockage levels, we postulate that the streptavidin pivots around its lone biotin-bound site, bending the end of the DNA fragment downward briefly. (Since the DNA is not anchored to a surface as in optical tweezers or AFM pulling experiments, this application of force appears to desorb the DNA from the nanopore wall without unzipping its strands or rupturing its bond with the streptavidin.) The deeper blockage depths of 42 to 50 nS observed during an event are attributed to the DNA fragment curling around the streptavidin until the flexural rigidity of the DNA fragment forces the streptavidin back toward the mouth of the pore. This process may be repeated as the MSA-DNA complex sticks and slides, effectively tumbling down the length of the pore before finally completing translocation. Although tumbling in nanopores has previously been reported for other proteins,(49, 129) we did not observe deep and prolonged multilevel events for streptavidin alone in the same pore (see Figure 2), emphasizing the role of the attached DNA.

The complex electrical signature profile of MSA bound to dsDNA at 200 mV is simplified at higher voltage, although the dwell times of detected events remain well above the resolution of our instrumentation. Increasing the applied voltage to 600 mV produces events such as those in Figure 5a, with a mean maximum blockage peak appearing at 27 ± 1 nS and an average dwell time of 100 ± 100 μs (Figure 5b). The 27 nS blockage depth is significantly deeper than that of streptavidin alone (7nS) at the same voltage and salt concentration (compare to Figure 2, blue), while the dwell time is actually shorter (100 μs versus 300 μs). The detected event rate doubles with the increase in voltage, although the most rapid events likely go undetected due to bandwidth limitations.

**Figure 5.**
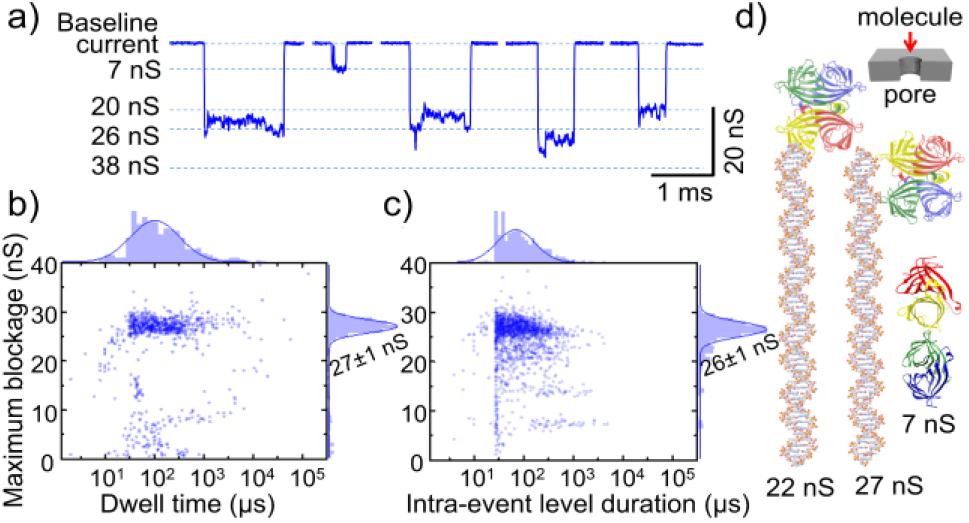
Nanopore detection of MSA-DNA complex in 4 M NaCl at 600 mV. Data obtained using the same pore as in Figures 1 and 4 (8.6 nm diameter, 11 nm thickness). a) Sample event traces. b) Overlay of dwell time and maximum blockage histograms. c) Overlay of intra-event level duration versus blockage level histograms for the same data. d) PDB-based cartoons of hypothesized MSA-DNA orientations. Inset shows orientation of depicted molecules relative to pore.

A closer look at typical events at 600 mV (Figure 5a) shows some intra-event variation in the current blockage, but nothing as discrete or as deep as the variations observed for the same system at 200 mV (Figure 4a). An overlay of the intra-event blockage level and level duration histograms shows a few events with maximum blockages around 15 nS and 7 nS, likely corresponding to trace amounts of free streptavidin in tetrameric or dimeric form (see Figure 2d for comparison), respectively, but the vast majority of blockage levels lie between 20 and 30 nS (Figure 5c).

The simple volume Equation 2 suggests the 27 ± 1 nS blockage produced by the protein-DNA complex at 600 mV corresponds to a volume of 150 ± 30 nm^3^, roughly 70% of the maximum volume of the MSA-DNA complex. Treating the MSA segment as a prolate spheroid with the electrical shape factor approach of Equation 3, we find values of 22 ± 5 nS and 30 ± 8 nS corresponding to the lengthwise or widthwise orientation of MSA, respectively (using dimensions of 4.5 x 5.8 nm and a biotin-bound volume of 133 nm^3^).

Once again, the resistors-in-series approach proves useful. By splitting the nanopore into cylindrical segments concentric with the nanopore axis, we can isolate the segment containing the MSA from the remainder of the pore. If both the DNA and MSA are treated as cylinders, the geometric model suggests that the protein is running lengthwise alongside the tail end of the dsDNA (27 ± 4 nS) or occasionally trailing lengthwise behind the DNA (22 ± 4 nS), as seen in Figure 5d. It is noteworthy that at 600 mV in the same pore, the MSA-DNA complex produces blockage depths ~20 nS greater than those of bare MSA (27 nS vs. 7 nS, respectively.) Since dsDNA fully occupying the pore length produces less than ~8 nS of blockage depth (see SI section 1), the disparity between the MSA-DNA blockage depth and that of the bare MSA is not fully explained by the additional volume of the DNA alone. Instead, the most probable explanation lies in the way biotin binding affects the structure of MSA. While an applied voltage of 600 mV induced a tetramer-dimer transition in the bare MSA, the same voltage leaves the tetramer intact once bound to biotinylated DNA, thus confirming reports of enhanced structural integrity conferred by biotin binding.(28)

To summarize our results for nanopore sensing of MSA bound to dsDNA, we find that discrete, significant fluctuations in the current amplitude at 200 mV correspond to conformational changes in the MSA-DNA complex, with the deepest blockages likely produced as the DNA fragment bends around the MSA. Although experimental data suggests that streptavidin alone ruptures at larger applied voltage (600 mV), the enhanced strength conferred to the streptavidin tetramer through its binding to biotinylated DNA evidently prevents its rupture on the timescale of its translocation through the pore. Further, the additional force at 600 mV (versus 200 mV) appears to drive the complex through the pore with far less non-specific adhesion to the pore surface, as the mean translocation time is shorter and the spread in measured blockage depths at 600 mV shows little sign of protein re-orientation or DNA bending.

The resistors-in-series approach offers a clear advantage for the study of multi-component assemblies. It allows consideration of the nanopore as a series of “slices,” with different shapes and volumes occupying each portion of the pore. Unlike existing approaches that would treat a protein-DNA binary system or multi-protein assembly as a spheroid or as an arbitrary volume (with no consideration for orientation effects), the resistors-in-series model provides a distribution of hypothesized states, with the most probable configuration identifiable through consideration of experimental forces and material properties (such as dipole moments).

## CONCLUSIONS

In conclusion, we have found that under an applied voltage of 600 mV in 4 M NaCl, the electric field within thin nanopores (~10 nm) likely destabilizes tetrameric MSA, rupturing the molecule into two dimers if protein-pore interactions provide sufficient opposing force. Under the same conditions, binding of MSA to biotinylated dsDNA appears to boost the MSA’s structural integrity and prevent the tetramer-dimer transition, although the complex is still stalled by strong interactions with the nanopore wall. The simple resistors-in-series approach described here allows stepwise analysis of real-time conformational changes of the protein-DNA complex in the nanopore, enabling the interpretation of complex electrical signatures observed. The insight provided by our approach suggests that DNA bends around the MSA at lower voltage, while at higher voltage the DNA remains rod-like with the MSA trailing behind.

The resistors-in-series approach could represent an important advancement in the analysis of biomolecules (e.g. proteins in various conformations) and biomolecular assemblies (e.g. protein-DNA complexes), allowing researchers to identify the range of hypothesized blockage depths associated with different configurations, orientations and folding states. Our model is applicable to a wide variety of biomaterials and can also be used in conjunction with existing models, since it allows sections of the pore to be treated individually. For example, the resistors-in-series approach allows us to combine the electrical shape factor model for a portion of the pore containing a spheroidal particle with the simple volume-based model for the remainder of the pore containing DNA. These results highlight the versatility of nanopore detection for a range of molecular assemblies and represent progress in the interpretation of complex translocation signals.

## AUTHOR CONTRIBUTIONS

A.C. performed research, analyzed data, and wrote the manuscript. V.T.C. edited the manuscript. Both authors have given approval to the final version of the manuscript.

## ACKNOWLEDGEMENTS

A. Carlsen would like to thank Z. Roelen for extensive discussion of the resistors-in-series model, M. Charron and K. Briggs for further discussion and critical review of the manuscript, and L. He for practical advice on the manuscript’s graphs and figures. The authors also express appreciation to M. Howarth and his group (including M. Jacobsen) for the generous and rapid delivery of monovalent streptavidin.

The authors would like to acknowledge the support of the Natural Sciences and Engineering Research Council of Canada (NSERC), grant CRDPJ 530554-2018.

## SUPPLEMENTAL INFORMATION

Supplemental information is available in the form of details of calculations (pore diameter, electrical shape factor, resistance-in-series model, diffusion coefficient, Stokes-Einstein radius, effective charge, effective confinement, capture radius) along with additional translocation data for MSA in KCl and LiCl, and controlled electromobility shift assay.

## REFERENCES

1. Laitinen, O.H., V.P. Hytönen, H.R. Nordlund, and M.S. Kulomaa. 2006. Genetically engineered avidins and streptavidins. Cell. Mol. Life Sci. 63:2992–3017.

2. Michael Green, N. 1990. Avidin and Streptavidin. Methods Enzymol. 184:51–67.

3. Wilchek, M., and E.A. Bayer. 1990. Introduction to avidin-biotin technology. Methods Enzymol. 184:5–13.

4. Li, H., S.H. Park, J.H. Reif, T.H. LaBean, and H. Yan. 2004. DNA-Templated Self-Assembly of Protein and Nanoparticle Linear Arrays. J. Am. Chem. Soc. 126:418–419.

5. Schultz, J., Y. Lin, J. Sanderson, Y. Zuo, D. Stone, R. Mallett, S. Wilbert, and D. Axworthy. 2000. A tetravalent single-chain antibody-streptavidin fusion protein for pretargeted lymphoma therapy. Cancer Res. 60:6663–9.

6. Neish, C.S., I.L. Martin, R.M. Henderson, and J.M. Edwardson. 2002. Direct visualization of ligand-protein interactions using atomic force microscopy. Br. J. Pharmacol. 135:1943–1950.

7. Jordan, C.E., A.G. Frutos, A.J. Thiel, and R.M. Corn. 1997. Surface Plasmon Resonance Imaging Measurements of DNA Hybridization Adsorption and Streptavidin/DNA Multilayer Formation at Chemically Modified Gold Surfaces. Anal. Chem. 69:4939–4947.

8. Grubmüller, H., B. Heymann, and P. Tavan. 1996. Ligand binding: molecular mechanics calculation of the streptavidin-biotin rupture force. Science. 271:997–9.

9. Weber, P.C., D.H. Ohlendorf, J.J. Wendoloski, and F.R. Salemme. 1989. Structural origins of high-affinity biotin binding to streptavidin. Science. 243:85–88.

10. Hendrickson, W.A., A. Pähler, J.L. Smith, Y. Satow, E.A. Merritt, and R.P. Phizackerley. 1989. Crystal structure of core streptavidin determined from multiwavelength anomalous diffraction of synchrotron radiation. Proc. Natl. Acad. Sci. U. S. A. 86:2190–4.

11. Carlsen, A.T., O.K. Zahid, J.A. Ruzicka, E.W. Taylor, and A.R. Hall. 2014. Selective detection and quantification of modified DNA with solid-state nanopores. Nano Lett. 14:5488–92.

12. Zahid, O.K., F. Wang, J.A. Ruzicka, E.W. Taylor, and A.R. Hall. 2016. Sequence-Specific Recognition of MicroRNAs and Other Short Nucleic Acids with Solid-State Nanopores. Nano Lett. 16:2033–2039.

13. Zahid, O.K., B.S. Zhao, C. He, and A.R. Hall. 2016. Quantifying mammalian genomic DNA hydroxymethylcytosine content using solid-state nanopores. Sci. Rep. 6:29565.

14. Wang, F., O.K. Zahid, B.E. Swain, D. Parsonage, T. Hollis, S. Harvey, F.W. Perrino, R.M. Kohli, E.W. Taylor, and A.R. Hall. 2017. Solid-State Nanopore Analysis of Diverse DNA Base Modifications Using a Modular Enzymatic Labeling Process. Nano Lett. 17:7110–7116.

15. Kong, J., N.A.W. Bell, and U.F. Keyser. 2016. Quantifying Nanomolar Protein Concentrations Using Designed DNA Carriers and Solid-State Nanopores. Nano Lett. 16:3557–3562.

16. Kong, J., J. Zhu, and U.F. Keyser. 2017. Single molecule based SNP detection using designed DNA carriers and solid-state nanopores. Chem. Commun. 53:436–439.

17. Chen, K., M. Juhasz, F. Gularek, E. Weinhold, Y. Tian, U.F. Keyser, and N.A.W. Bell. 2017. Ionic Current-Based Mapping of Short Sequence Motifs in Single DNA Molecules Using Solid-State Nanopores. Nano Lett. 17:5199–5205.

18. Hu, R., J. V. Rodrigues, P. Waduge, H. Yamazaki, B. Cressiot, Y. Chishti, L. Makowski, D. Yu, E. Shakhnovich, Q. Zhao, and M. Wanunu. 2018. Differential Enzyme Flexibility Probed Using Solid-State Nanopores. ACS Nano. 12:4494–4502.

19. Yusko, E.C., B.R. Bruhn, O.M. Eggenberger, J. Houghtaling, R.C. Rollings, N.C. Walsh, S. Nandivada, M. Pindrus, A.R. Hall, D. Sept, J. Li, D.S. Kalonia, and M. Mayer. 2017. Real-time shape approximation and fingerprinting of single proteins using a nanopore. Nat. Nanotechnol. 12:360–367.

20. Briggs, K., M. Charron, H. Kwok, T. Le, S. Chahal, J. Bustamante, M. Waugh, and V. Tabard-Cossa. 2015. Kinetics of nanopore fabrication during controlled breakdown of dielectric membranes in solution. Nanotechnology. 26:084004.

21. Kwok, H., K. Briggs, and V. Tabard-Cossa. 2014. Nanopore fabrication by controlled dielectric breakdown. PLoS One. 9:e92880.

22. Waugh, M., K. Briggs, D. Gunn, M. Gibeault, S. King, Q. Ingram, A.M. Jimenez, S. Berryman, D. Lomovtsev, L. Andrzejewski, and V. Tabard-Cossa. 2020. Solid-state nanopore fabrication by automated controlled breakdown. Nat. Protoc. 15:122–143.

23. Forstater, J.H., K. Briggs, J.W.F. Robertson, J. Ettedgui, O. Marie-Rose, C. Vaz, J.J. Kasianowicz, V. Tabard-Cossa, and A. Balijepalli. 2016. MOSAIC: A modular single-molecule analysis interface for decoding multistate nanopore data. Anal. Chem. 88:11900–11907.

24. Howarth, M., D.J.-F. Chinnapen, K. Gerrow, P.C. Dorrestein, M.R. Grandy, N.L. Kelleher, A. El-Husseini, and A.Y. Ting. 2006. A monovalent streptavidin with a single femtomolar biotin binding site. Nat. Methods. 3:267–273.

25. Fairhead, M., D. Krndija, E.D. Lowe, and M. Howarth. 2014. Plug-and-Play Pairing via Defined Divalent Streptavidins. J. Mol. Biol. 426:199–214.

26. Moreland, J.L., A. Gramada, O. V Buzko, Q. Zhang, and P.E. Bourne. 2005. The Molecular Biology Toolkit (MBT): a modular platform for developing molecular visualization applications. BMC Bioinformatics. 6:21.

27. Wu, S.-C., and S.-L. Wong. 2005. Engineering Soluble Monomeric Streptavidin with Reversible Biotin Binding Capability. J. Biol. Chem. 280:23225–23231.

28. Kurzban, G.P., E.A. Bayer, M. Wilchek, and P.M. Horowitz. 1991. The quaternary structure of streptavidin in urea. J. Biol. Chem. 266:14470–7.

29. Kim, M., C.-C. Wang, F. Benedetti, M. Rabbi, V. Bennett, and P.E. Marszalek. 2011. Nanomechanics of streptavidin hubs for molecular materials. Adv. Mater. 23:5684–8.

30. Wu, S.-C., and S.-L. Wong. 2005. Engineering soluble monomeric streptavidin with reversible biotin binding capability. J. Biol. Chem. 280:23225–31.

31. Sivasankar, S., S. Subramaniam, and D. Leckband. 1998. Direct molecular level measurements of the electrostatic properties of a protein surface. Proc. Natl. Acad. Sci. U. S. A. 95:12961–6.

32. Innovagen. 2014. Peptide calculator. https://pepcalc.com/. 2019-09-10.

33. Kowalczyk, S.W., A.Y. Grosberg, Y. Rabin, and C. Dekker. 2011. Modeling the conductance and DNA blockade of solid-state nanopores. Nanotechnology. 22:315101.

34. Golibersuch, D.C. 1973. Observation of aspherical particle rotation in Poiseuille flow via the resistance pulse technique. I. Application to human erythrocytes. Biophys. J. 13:265–80.

35. Qin, Z., J. Zhe, and G.-X. Wang. 2011. Effects of particle’s off-axis position, shape, orientation and entry position on resistance changes of micro Coulter counting devices. Meas. Sci. Technol. 22:045804.

36. Yusko, E.C., J.M. Johnson, S. Majd, P. Prangkio, R.C. Rollings, J. Li, J. Yang, and M. Mayer. 2011. Controlling protein translocation through nanopores with bio-inspired fluid walls. Nat. Nanotechnol. 6:253–60.

37. DeBlois, R.W., and C.P. Bean. 1970. Counting and sizing of submicron particles by the resistive pulse technique. Rev. Sci. Instrum. 41:909–916.

38. Fologea, D., B. Ledden, D.S. McNabb, and J. Li. 2007. Electrical characterization of protein molecules by a solid-state nanopore. Appl. Phys. Lett. 91:053901.

39. Ledden, B., D. Fologea, D.S. Talaga, and J. Li. 2011. Sensing Single Protein Molecules with Solid-State Nanopores. In: Nanopores. Springer US. pp. 129–150.

40. Hall, J.E. 1975. Access resistance of a small circular pore. J. Gen. Physiol. 66:531–2.

41. Talaga, D.S., and J. Li. 2009. Single-molecule protein unfolding in solid state nanopores. J. Am. Chem. Soc. 131:9287–97.

42. Houghtaling, J., C. Ying, O.M. Eggenberger, A. Fennouri, S. Nandivada, M. Acharjee, J. Li, A.R. Hall, and M. Mayer. 2019. Estimation of Shape, Volume, and Dipole Moment of Individual Proteins Freely Transiting a Synthetic Nanopore. ACS Nano. 13:5231–5242.

43. Freedman, K.J., S.R. Haq, J.B. Edel, P. Jemth, and M.J. Kim. 2013. Single molecule unfolding and stretching of protein domains inside a solid-state nanopore by electric field. Sci. Rep. 3:1638.

44. Häussling, L., B. Michel, H. Ringsdorf, and H. Rohrer. 1991. Direct Observation of Streptavidin Specifically Adsorbed on Biotin-Functionalized Self-Assembled Monolayers with the Scanning Tunneling Microscope. Angew. Chemie Int. Ed. English. 30:569–572.

45. Niedzwiecki, D.J., J. Grazul, and L. Movileanu. 2010. Single-molecule observation of protein adsorption onto an inorganic surface. J. Am. Chem. Soc. 132:10816–10822.

46. Williams, E.H., A. V Davydov, A. Motayed, S.G. Sundaresan, P. Bocchini, L.J. Richter, G. Stan, K. Steffens, R. Zangmeister, J.A. Schreifels, and V. Rao. 2012. Immobilization of streptavidin on 4H-SiC for biosensor development. Appl. Surf. Sci. 258:6056–6063.

47. Spinke, J., M. Liley, F.-J. Schmitt, H.-J. Guder, L. Angermaier, and W. Knoll. 1993. Molecular recognition at self-assembled monolayers: Optimization of surface functionalization. J. Chem. Phys. 99:7012.

48. Varongchayakul, N., D. Huttner, M.W. Grinstaff, and A. Meller. 2018. Sensing Native Protein Solution Structures Using a Solid-state Nanopore: Unraveling the States of VEGF. Sci. Rep. 8:1017.

49. Varongchayakul, N., J. Song, A. Meller, and M.W. Grinstaff. 2018. Single-molecule protein sensing in a nanopore: a tutorial. Chem. Soc. Rev. 47:8512–8524.

50. Van Oss, C.J., R.F. Giese, P.M. Bronson, A. Docoslis, P. Edwards, and W.T. Ruyechan. 2003. Macroscopic-scale surface properties of streptavidin and their influence on aspecific interactions between streptavidin and dissolved biopolymers. Colloids Surfaces B Biointerfaces. 30:25–36.

51. Carlsen, A.T., O.K. Zahid, J. Ruzicka, E.W. Taylor, and A.R. Hall. 2014. Interpreting the Conductance Blockades of DNA Translocations through Solid-State Nanopores. ACS Nano. 8:4754–4760.

52. He, L.-Z., and B. Niemeyer. 2003. A Novel Correlation for Protein Diffusion Coefficients Based on Molecular Weight and Radius of Gyration. Biotechnol. Prog. 19:544–548.

53. Muthukumar, M. 2014. Communication: Charge, diffusion, and mobility of proteins through nanopores. J. Chem. Phys. 141:081104.

54. Larkin, J., R.Y. Henley, M. Muthukumar, J.K. Rosenstein, and M. Wanunu. 2014. High-Bandwidth Protein Analysis Using Solid-State Nanopores. Biophys. J. 106:696–704.

55. Plesa, C., S.W. Kowalczyk, R. Zinsmeester, A.Y. Grosberg, Y. Rabin, and C. Dekker. 2013. Fast translocation of proteins through solid state nanopores. Nano Lett. 13:658–663.

56. Haridasan, N., S.K. Kannam, S. Mogurampelly, and S.P. Sathian. 2018. Translational mobilities of proteins in nanochannels: A coarse-grained molecular dynamics study. Phys. Rev. E. 97:062415.

57. Wanunu, M., J. Sutin, B. McNally, A. Chow, and A. Meller. 2008. DNA Translocation Governed by Interactions with Solid-State Nanopores. Biophys. J. 95:4716–4725.

58. Bowen, W.R., A.W. Mohammad, and N. Hilal. 1997. Characterisation of nanofiltration membranes for predictive purposes — use of salts, uncharged solutes and atomic force microscopy. J. Memb. Sci. 126:91–105.

59. Kannam, S.K., and M.T. Downton. 2017. Translational diffusion of proteins in nanochannels. J. Chem. Phys. 146:054108.

60. Berg, H.C. 2018. Random Walks in Biology. Princeton University Press.

61. Waduge, P., R. Hu, P. Bandarkar, H. Yamazaki, B. Cressiot, Q. Zhao, P.C. Whitford, and M. Wanunu. 2017. Nanopore-Based Measurements of Protein Size, Fluctuations, and Conformational Changes. ACS Nano. 11:5706–5716.

62. Nakane, J., M. Akeson, and A. Marziali. 2002. Evaluation of nanopores as candidates for electronic analyte detection. Electrophoresis. 23:2592–2601.

63. Qiao, L., M. Ignacio, and G.W. Slater. 2019. Voltage-driven translocation: Defining a capture radius. J. Chem. Phys. 151:244902.

64. Wanunu, M., W. Morrison, Y. Rabin, A.Y. Grosberg, and A. Meller. 2010. Electrostatic focusing of unlabelled DNA into nanoscale pores using a salt gradient. Nat. Nanotechnol. 5:160–165.

65. Grosberg, A.Y., and Y. Rabin. 2010. DNA capture into a nanopore: Interplay of diffusion and electrohydrodynamics. J. Chem. Phys. 133:165102.

66. Muthukumar, M. 2014. Macromolecular Mechanisms of Protein Translocation. Protein Pept. Lett. 21:209–216.

67. Cressiot, B., A. Oukhaled, G. Patriarche, M. Pastoriza-Gallego, J.-M. Betton, L. Auvray, M. Muthukumar, L. Bacri, and J. Pelta. 2012. Protein Transport through a Narrow Solid-State Nanopore at High Voltage: Experiments and Theory. ACS Nano. 6:6236–6243.

68. Oukhaled, A., B. Cressiot, L. Bacri, M. Pastoriza-Gallego, J.-M. Betton, E. Bourhis, R. Jede, J. Gierak, L. Auvray, and J. Pelta. 2011. Dynamics of Completely Unfolded and Native Proteins through Solid-State Nanopores as a Function of Electric Driving Force. ACS Nano. 5:3628–3638.

69. Freedman, K.J., M. Jürgens, A. Prabhu, C.W. Ahn, P. Jemth, J.B. Edel, and M.J. Kim. 2011. Chemical, thermal, and electric field induced unfolding of single protein molecules studied using nanopores. Anal. Chem. 83:5137–5144.

70. Ojeda-May, P., and M.E. Garcia. 2010. Electric field-driven disruption of a native beta-sheet protein conformation and generation of a helix-structure. Biophys. J. 99:595–9.

71. Bekard, I., and D.E. Dunstan. 2014. Electric field induced changes in protein conformation. Soft Matter. 10:431–437.

72. Rochu, D., T. Pernet, F. Renault, C. Bon, and P. Masson. 2001. Dual effect of high electric field in capillary electrophoresis study of the conformational stability of Bungarus fasciatus acetylcholinesterase. J. Chromatogr. A. 910:347–357.

73. Astrakas, L., C. Gousias, and M. Tzaphlidou. 2011. Electric field effects on chignolin conformation. J. Appl. Phys. 109:094702.

74. Jiang, Z., L. You, W. Dou, T. Sun, and P. Xu. 2019. Effects of an electric field on the conformational transition of the protein: A molecular dynamics simulation study. Polymers (Basel). 11:282.

75. Waner, M.J., I. Navrotskaya, A. Bain, E.D. Oldham, and D.P. Mascotti. 2004. Thermal and sodium dodecylsulfate induced transitions of streptavidin. Biophys. J. 87:2701–2713.

76. Kim, M., C.-C. Wang, F. Benedetti, M. Rabbi, V. Bennett, and P.E. Marszalek. 2011. Nanomechanics of streptavidin hubs for molecular materials. Adv. Mater. 23:5684–8.

77. Kannam, S.K., S.C. Kim, P.R. Rogers, N. Gunn, J. Wagner, S. Harrer, and M.T. Downton. 2014. Sensing of protein molecules through nanopores: a molecular dynamics study. Nanotechnology. 25:155502.

78. Wang, X., Y. Li, X. He, S. Chen, and J.Z.H. Zhang. 2014. Effect of strong electric field on the conformational integrity of insulin. J. Phys. Chem. A. 118:8942–8952.

79. Firnkes, M., D. Pedone, J. Knezevic, M. Döblinger, and U. Rant. 2010. Electrically Facilitated Translocations of Proteins through Silicon Nitride Nanopores: Conjoint and Competitive Action of Diffusion, Electrophoresis, and Electroosmosis. Nano Lett. 10:2162–2167.

80. English, N.J., G.Y. Solomentsev, and P. O’Brien. 2009. Nonequilibrium molecular dynamics study of electric and low-frequency microwave fields on hen egg white lysozyme. J. Chem. Phys. 131:035106.

81. Felder, C.E., J. Prilusky, I. Silman, and J.L. Sussman. 2007. A server and database for dipole moments of proteins. Nucleic Acids Res. 35:W512.

82. Weizmann Institute. 2007. Protein Dipole Moments Server. https://dipole.weizmann.ac.il/dipol/. 2019-09-18.

83. Awsiuk, K., P. Petrou, A. Thanassoulas, and J. Raczkowska. 2019. Orientation of Biotin-Binding Sites in Streptavidin Adsorbed onto the Surface of Polythiophene Films. Langmuir. 35:3058–3066.

84. Squires, A., E. Atas, and A. Meller. 2015. Nanopore sensing of individual transcription factors bound to DNA. Sci. Rep. 5:11643.

85. Dudko, O.K., J. Mathé, A. Szabo, A. Meller, and G. Hummer. 2007. Extracting kinetics from single-molecule force spectroscopy: nanopore unzipping of DNA hairpins. Biophys. J. 92:4188–95.

86. Comer, J., A. Ho, and A. Aksimentiev. 2012. Toward detection of DNA-bound proteins using solid-state nanopores: Insights from computer simulations. Electrophoresis. 33:3466–3479.

87. Keyser, U.F., B.N. Koeleman, S. van Dorp, D. Krapf, R.M.M. Smeets, S.G. Lemay, N.H. Dekker, and C. Dekker. 2006. Direct force measurements on DNA in a solid-state nanopore. Nat. Phys. 2:473–477.

88. Spiering, A., S. Getfert, A. Sischka, P. Reimann, and D. Anselmetti. 2011. Nanopore translocation dynamics of a single DNA-bound protein. Nano Lett. 11:2978–2982.

89. Tabard-Cossa, V., M. Wiggin, D. Trivedi, N.N. Jetha, J.R. Dwyer, and A. Marziali. 2009. Singlemolecule bonds characterized by solid-state nanopore force spectroscopy. ACS Nano. 3:3009–14.

90. Lindman, S., W.F. Xue, O. Szczepankiewicz, M.C. Bauer, H. Nilsson, and S. Linse. 2006. Salting the charged surface: pH and salt dependence of protein G B1 stability. Biophys. J. 90:2911–2921.

91. Perez-Jimenez, R., R. Godoy-Ruiz, B. Ibarra-Molero, and J.M. Sanchez-Ruiz. 2004. The Efficiency of Different Salts to Screen Charge Interactions in Proteins: A Hofmeister Effect? Biophys. J. 86:2414–2429.

92. van Dorp, S., U.F. Keyser, N.H. Dekker, C. Dekker, and S.G. Lemay. 2009. Origin of the electrophoretic force on DNA in solid-state nanopores. Nat. Phys. 5:347–351.

93. Luan, B., T. Huynh, J. Li, and R. Zhou. 2016. Nanomechanics of protein unfolding outside a generic nanopore. ACS Nano. 10:317–323.

94. Toschi, F., F. Lugli, F. Biscarini, and F. Zerbetto. 2009. Effects of Electric Field Stress on a β-Amyloid Peptide. J. Phys. Chem. B. 113:369–376.

95. Marracino, P., F. Apollonio, M. Liberti, G. d’Inzeo, and A. Amadei. 2013. Effect of High Exogenous Electric Pulses on Protein Conformation: Myoglobin as a Case Study. J. Phys. Chem. B. 117:2273–2279.

96. Okur, H.I., J. Hladílková, K.B. Rembert, Y. Cho, J. Heyda, J. Dzubiella, P.S. Cremer, and P. Jungwirth. 2017. Beyond the Hofmeister Series: Ion-Specific Effects on Proteins and Their Biological Functions. J. Phys. Chem. B. 121:1997–2014.

97. Arakawa, T., and S.N. Timasheff. 1982. Preferential Interactions of Proteins with Salts in Concentrated Solutions. Biochemistry. 21:6545–6552.

98. Baldwin, R.L. 1996. How Hofmeister ion interactions affect protein stability. Biophys. J. 71:2056–2063.

99. Gronau, G., Z. Qin, and M.J. Buehler. 2013. Effect of sodium chloride on the structure and stability of spider silk’s N-terminal protein domain. Biomater. Sci. 1:276–284.

100. Purohit, A., J.K. England, L.G. Douma, F. Tondnevis, L.B. Bloom, and M. Levitus. 2017. Electrostatic Interactions at the Dimer Interface Stabilize the E. coli β Sliding Clamp. Biophys. J. 113:794–804.

101. Shima, S., C. Tziatzios, D. Schubert, H. Fukada, K. Takahashi, U. Ermler, and R.K. Thauer. 1998. Lyotropic-salt-induced changes in monomer/dimer/tetramer association equilibrium of formyltransferase from the hyperthermophilic Methanopyrus kandleri in relation to the activity and thermostability of the enzyme. Eur. J. Biochem. 258:85–92.

102. Von Hippel, P.H., and K.Y. Wong. 1965. On the conformational stability of globular proteins. The effects of various electrolytes and nonelectrolytes on the thermal ribonuclease transition. J. Biol. Chem. 240:3909–3923.

103. Collins, K.D., and M.W. Washabaugh. 1985. The Hofmeister effect and the behaviour of water at interfaces. Q. Rev. Biophys. 18:323–422.

104. Zhang, Y., and P.S. Cremer. 2006. Interactions between macromolecules and ions: the Hofmeister series. Curr. Opin. Chem. Biol. 10:658–663.

105. Hofmeister, F. 1888. Zur Lehre von der Wirkung der Salze – Zweite Mittheilung. Arch. für Exp. Pathol. und Pharmakologie. 24:247–260.

106. Kunz, W., J. Henle, and B.W. Ninham. 2004. “Zur Lehre von der Wirkung der Salze” (about the science of the effect of salts): Franz Hofmeister’s historical papers. In: Current Opinion in Colloid and Interface Science. Elsevier. pp. 19–37.

107. Gibb, B.C. 2019. Hofmeister’s curse. Nat. Chem. 11:963–965.

108. Schwierz, N., D. Horinek, U. Sivan, and R.R. Netz. 2016. Reversed Hofmeister series—The rule rather than the exception. Curr. Opin. Colloid Interface Sci. 23:10–18.

109. Gibb, C.L.D., and B.C. Gibb. 2011. Anion binding to hydrophobic concavity is central to the salting-in effects of hofmeister chaotropes. J. Am. Chem. Soc. 133:7344–7347.

110. Grover, P.K., and R.L. Ryall. 2005. Critical appraisal of salting-out and its implications for chemical and biological sciences. Chem. Rev. 105:1–10.

111. Hyde, A.M., S.L. Zultanski, J.H. Waldman, Y.L. Zhong, M. Shevlin, and F. Peng. 2017. General Principles and Strategies for Salting-Out Informed by the Hofmeister Series. Org. Process Res. Dev. 21:1355–1370.

112. Cao, Q. 2016. Hofmeister effect for electrokinetic transport at ordered DNA layers. Microfluid. Nanofluidics. 20:1–10.

113. Tsumoto, K., D. Ejima, A.M. Senczuk, Y. Kita, and T. Arakawa. 2007. Effects of salts on protein–surface interactions: applications for column chromatography. J. Pharm. Sci. 96:1677–1690.

114. Le Trong, I., Z. Wang, D.E. Hyre, T.P. Lybrand, P.S. Stayton, and R.E. Stenkamp. 2011. Streptavidin and its biotin complex at atomic resolution. Acta Crystallogr. D. Biol. Crystallogr. 67:813–21.

115. D’Agata, R., P. Palladino, and G. Spoto. 2017. Streptavidin-coated gold nanoparticles: Critical role of oligonucleotides on stability and fractal aggregation. Beilstein J. Nanotechnol. 8:1–11.

116. Karau, P., and V. Tabard-Cossa. 2018. Capture and Translocation Characteristics of Short Branched DNA Labels in Solid-State Nanopores. ACS Sensors. 3:1308–1315.

117. Shi, X., Q. Li, R. Gao, W. Si, S.-C. Liu, A. Aksimentiev, and Y.-T. Long. 2018. Dynamics of a Molecular Plug Docked onto a Solid-State Nanopore. J. Phys. Chem. Lett. 9:4686–4694.

118. Gross, P., N. Laurens, L.B. Oddershede, U. Bockelmann, E.J.G. Peterman, and G.J.L. Wuite. 2011. Quantifying how DNA stretches, melts and changes twist under tension. Nat. Phys. 7:731–736.

119. Bustamante, C., J.F. Marko, E.D. Siggia, and S. Smith. 1994. Entropic elasticity of λ-phage DNA. Science (80-.). 265:1599–1600.

120. Marko, J.F., and E.D. Siggia. 1995. Stretching DNA. Macromolecules. 28:8759–8770.

121. Hagerman, P.J. 1988. Flexibility of DNA. Annu. Rev. Biophys. Biophys. Chem. 17:265–286.

122. Zeida, A., M.R. Machado, P.D. Dans, and S. Pantano. 2012. Breathing, bubbling, and bending: DNA flexibility from multimicrosecond simulations. Phys. Rev. E. 86:021903.

123. Yuan, C., H. Chen, X.W. Lou, and L.A. Archer. 2008. DNA Bending Stiffness on Small Length Scales. Phys. Rev. Lett. 100:018102.

124. Garcia, H.G., P. Grayson, L. Han, M. Inamdar, J. Kondev, P.C. Nelson, R. Phillips, J. Widom, and P.A. Wiggins. 2007. Biological consequences of tightly bent DNA: the other life of a macromolecular celebrity. Biopolymers. 85:115–30.

125. David Boal. 2002. Mechanics of the Cell. New York: Cambridge University Press.

126. Savelyev, A. 2012. Do monovalent mobile ions affect DNA’s flexibility at high salt content? Phys. Chem. Chem. Phys. 14:2250–2254.

127. Kriegel, F., N. Ermann, R. Forbes, D. Dulin, N.H. Dekker, and J. Lipfert. 2017. Probing the salt dependence of the torsional stiffness of DNA by multiplexed magnetic torque tweezers. Nucleic Acids Res. 45:5920–5929.

128. Cressiot, B., A. Oukhaled, L. Bacri, and J. Pelta. 2014. Focus on Protein Unfolding Through Nanopores. Bionanoscience. 4:111–118.

129. Zernia, S., N.J. Van Der Heide, N.S. Galenkamp, G. Gouridis, and G. Maglia. 2020. Current Blockades of Proteins inside Nanopores for Real-Time Metabolome Analysis. ACS Nano. 14:2296–2307.

