## Supporting Information for "Mapping shifts in nanopore signal to changes in protein and protein-DNA conformation"

Autumn T. Carlsen, Vincent Tabard Cossa

Department of Physics, University of Ottawa, Ottawa, Ontario K1N 6N5, Canada

#### Supplemental Information Table of Contents

S1. Methods of pore diameter calculation and error propagation

S2. Electromobility Shift Assay (EMSA)

S3. Calculation of electrical shape factor

S4. Calculation of resistors-in-series model for a single macromolecule within the pore

S5. Calculation of diffusion coefficient

S6. Calculation of Stokes-Einstein radius

S7. Calculation of effective charge

S8. Effect of confinement

S9. Comparison of MSA translocation at applied voltages of 200 mV versus 800 mV through the same pore

S10. Calculation of capture radius

S11. Discrete intra-event levels in KCl and LiCl at 200 mV

S12. Calculation of resistors-in-series model for two macromolecules

#### S1. Methods of pore diameter calculation and error propagation

To validate pore performance and determine pore geometry, we began each experimental set by translocating 1 kbp DNA. Figure S1 shows a sample current trace along with the observed distribution of the maximum current blockage in 4 M NaCl at 200 mV for the pore used to acquire the data in Figures 2, 4 and 5 of the main manuscript.

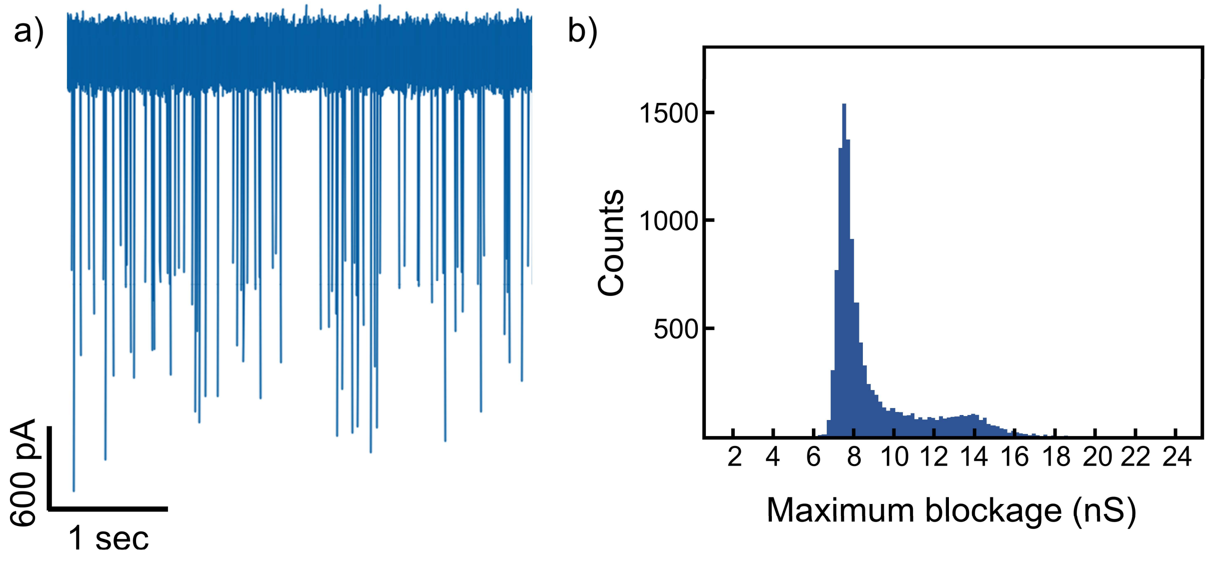


**Figure S1 – Nanopore detection of double-stranded 1 kbp DNA in 4 M NaCl at 200 mV.** a) Typical 4-second current trace for translocations of 1 kbp dsDNA through the solid-state nanopore subsequently used to collect MSA and MSA-DNA events (data presented in Figures 2, 4 and 5 of the main manuscript). b) Maximum blockage histogram for 11,548 events. Analysis of these DNA translocations revealed the pore to be 8.4 nm in diameter and 11 nm thick.

Analysis of over 11,500 events revealed a clear peak in maximum blockage depth corresponding to unfolded 1 kbp DNA at 7.6 ± 0.4 nS. (Translocation of partially folded 1 kbp DNA produced a small, secondary peak at nearly twice the unfolded value.) The maximum blockage depth obtained for unfolded DNA was used to calibrate the effective thickness and diameter for each pore used in this work. The DNA-calibrated thickness of the membrane ($H_{eff}$ = 11 nm) was calculated using the simplest conductance model for a cylindrical pore:(1)

$$\Delta G_{DNA} = \frac{\sigma\pi d_{DNA}^2}{4H_{eff}} = \frac{\sigma(\pi d_{DNA}^2 H_{eff}/4)}{H_{eff}^2} = \frac{\sigma \Lambda_{DNA}}{H_{eff}^2} \qquad (1)$$

by substituting the crystallographic edge-to-edge distance of 2.2 nm for the diameter of double-stranded DNA ($d_{DNA}$), along with the measured solution conductivity ($\sigma = 22\ S/m$) and the first-level blockage depth ($\Delta G_{DNA} = 7.6\ nS$). Note that the final term in the equation includes DNA volume ($\Lambda_{DNA} = \pi d_{DNA}^2 H_{eff}/4$) to reproduce a form often found in literature.

For the translocation of double-stranded DNA in narrow pores ($d_p < 15\ nm$), Eq. 1 provides a reasonable approximation of the pore length, since the volume fraction of the DNA within the pore is much larger than the volume fraction outside,(1) meaning:

$$\Delta G_{DNA} = G_0 - G_{pore\ with\ DNA} \cong \frac{\sigma\pi d_p^2}{4H_{eff}} - \frac{\sigma\pi(d_p^2 - d_{DNA}^2)}{4H_{eff}} = \frac{\sigma\pi d_{DNA}^2}{4H_{eff}}. \tag{2}$$

where $G_0$ is the calculated open pore conductance and $G_{pore\ with\ DNA}$ is the calculated conductance of the pore while its cross-sectional area is effectively reduced by the presence of DNA. To determine the effective pore diameter, we substitute the DNA-calibrated length (11 nm) of Eq. 1 and the measured open pore conductance ($G_{open\ pore}$ = 69 nS) into a conductance model incorporating access region effects as follows:

$$G_{open\ pore} = \sigma\left(\frac{4H_{eff}}{\pi d_p^2} + \frac{1}{d_p}\right)^{-1} = \sigma\left(\frac{\pi d_p^2}{4H_{eff} + \pi d_p}\right) \cong \sigma\left(\frac{\pi d_p^2/4}{H_{eff} + 0.8 d_p}\right). \tag{3}$$

Using experimentally determined values for the effective pore diameter ($d_p$ = 8.4 nm) and DNA-calibrated pore length ($H_{eff}$ = 11 nm), we obtain the correct blockage depth for DNA using Eq. 2.

We use the same values for pore length (11 nm) and pore diameter (8.4 nm) in calculations for the protein and protein-DNA complex in all models described in this paper. Errors are calculated using standard error propagation techniques with the following assumptions:

- Uncertainty in pore thickness = ± 1 nm (e.g. 11 ± 1 nm)
- Uncertainty in pore diameter = ± 0.2 nm (e.g. 8.4 ± 0.2 nm)
- Uncertainty in molecular axes (length, diameter) = ± 0.2 nm (e.g. 4.2 ± 0.2 nm)
- Uncertainty in conductivity = ± 0.2 S/m (e.g. 22 ± 0.2 S/m)
- Uncertainty in baseline current (in electrical shape factor model) = ± 3% (e.g. 71 ± 2 nS)
- Uncertainty in analyte molar concentration = 10% (e.g. 30 ± 3 nM).

#### S2. Electromobility Shift Assay (EMSA)

The quality of the MSA-DNA complex was confirmed by multiple gel electrophoresis experiments. In the example gel below (Figure S2), MSA was incubated with 56 bp biotinylated DNA in 0.3X PBS buffer for one hour at room temperature at molar ratios ranging from 0:1 to 3:1 (MSA to DNA). Samples were subsequently loaded onto a 2% agarose gel with GelRed nucleic acid stain for visualization.

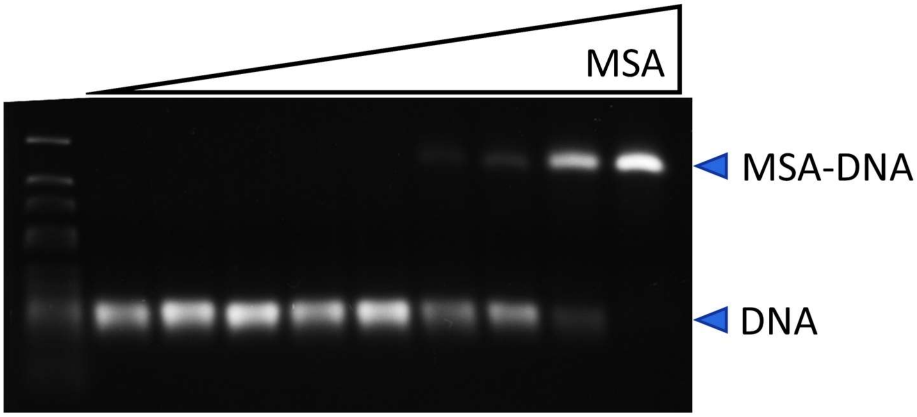


**Figure S2 – Example gel confirming quality of MSA-DNA complexes.** All lanes (excluding the Generuler Ultra Low Range DNA ladder in lane 1 on the far left) contain the same amount of biotinylated DNA (21 ng). Lanes 3 through 10 include increasing amount of streptavidin with molar concentration ranging from 0.3 times to 3.1 times the molar concentration of DNA (60 nM). From left to right, the lanes contain ladder, 0x, 0.3x, 0.7x, 1x, 1.3x, 1.5x, 1.8x, 2.2x and 3.1x streptavidin to biotinylated DNA.

### S3. Calculation of electrical shape factor

According to Grover,(2) Golibersuch,(3) Yusko(4) and Houghtaling et al.,(5) the orientation of a spheroid will generate a range of electrical shape factors $\gamma$ with $\gamma_{para}$ and $\gamma_{perp}$ produced when the spheroid's axis of symmetry or revolution (equivalent to the length value here) lies parallel or perpendicular to the electric field, respectively. For a prolate spheroid ($m$ > 1, where $m$ is the ratio of length $a$ to diameter $b$), $\gamma_{perp}$ will correspond to the maximum blockage depth ($\Delta I/I_0$), produced when the longer axis (length) of the spheroid is perpendicular to the electric field. The minimum blockage will be produced when the spheroid's longer axis is parallel to the electric field, corresponding to $\gamma_{para}$.

Following Yusko et al.(4) closely, the electrical shape factors ($\gamma_{para}$ and $\gamma_{perp}$) can be obtained from the depolarization factors $\eta_{para}$ and $\eta_{perp}$ as follows:

$$\gamma_{para} = \frac{1}{1-\eta_{para}} \; ; \; \gamma_{perp} = \frac{1}{1-\eta_{perp}} \tag{4}$$

where $\eta_{para}$ for a prolate spheroid (m > 1, or length > diameter) is defined as:

$$\eta_{para} = \frac{1}{m^2-1}\left(\frac{m}{\sqrt{m^2-1}}\ln\left(m+\sqrt{m^2-1}\right)-1\right) \tag{5}$$

and $\eta_{para}$ for an oblate spheroid (m<1 or length <diameter) is defined as:

$$\eta_{para} = \frac{1}{1-m^2}\left(1-\frac{m}{\sqrt{1-m^2}}\cos^{-1}(\mathrm{m})\right). \tag{6}$$

For both prolate and oblate cases, $\eta_{perp}$ is defined as $\eta_{perp} = (1-\eta_{para})/2$.

#### S4. Calculation of resistors-in-series model for a single macromolecule within the pore

The resistors-in-series model treats the nanopore as a cylinder with multiple segments concentric with the nanopore's central axis. When a portion of the pore is occupied as seen in Figure S3, the total resistance of the pore with the molecule inside ($R_{pore\ with\ molecule}$) can be obtained by adding the resistance of the occupied portion of the pore ($R_{occupied\ portion}$) with the resistance of the empty portion of the pore ($R_{empty\ portion}$) as follows:

$$R_{pore\ with\ molecule} = R_{empty\ portion} + R_{occupied\ portion} \quad (7)$$

$$= \frac{1}{\sigma}\frac{(H_{eff}-l_m)}{A_p} + \frac{1}{\sigma}\frac{l_m}{A_p-A_m} = \frac{1}{\sigma}\frac{A_pH_{eff}-A_m(H_{eff}-l_m)}{A_p(A_p-A_m)}$$

where $\sigma$ is the solution conductivity, $H_{eff}$ is the effective pore length determined by DNA blockage depth (see section S1), $l_m$ is the molecule length, $A_p$ is the cross-sectional area of the pore, and $A_m$ is the cross-sectional area of the molecule.

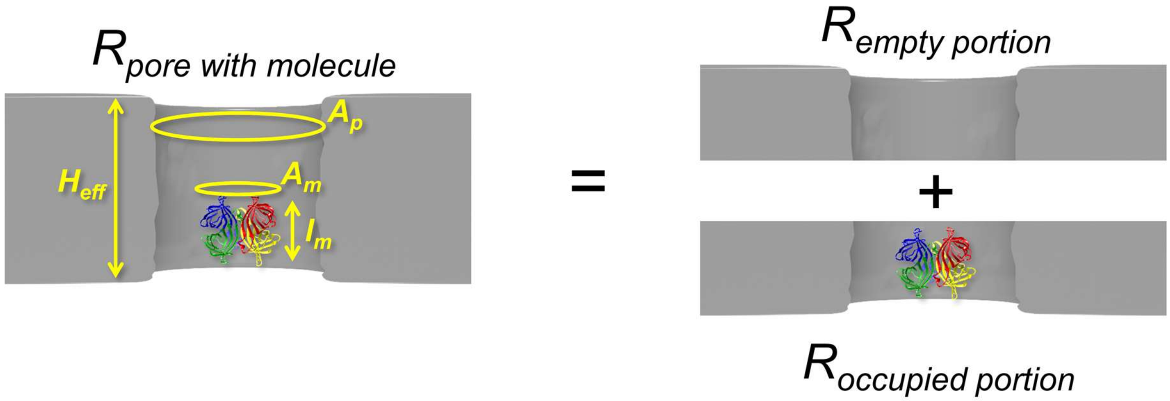


**Figure S3 – Visual diagram demonstrating the equivalence the resistance of a pore with a molecule to the resistance of two segments of the pore in series.** The area of the pore $A_p$, effective length of the pore $H_{eff}$, area of the molecule $A_m$ and length of the molecule $l_m$ are denoted.

Since conductance equals the reciprocal of resistance ($G = R^{-1}$), we find the conductance of the pore with the molecule to be:

$$R^{-1}_{pore\ with\ molecule} = G_{pore\ with\ molecule} = \frac{\sigma A_p(A_p-A_m)}{A_pH_{eff}-A_m(H_{eff}-l_m)}. \quad (8)$$

The change in conductance ($\Delta G$) due to translocation of the molecule can then be expressed as:

$$\Delta G = G_0 - G_{pore\ with\ molecule} = \frac{\sigma A_p}{H_{eff}} - \frac{\sigma A_p(A_p-A_m)}{A_pH_{eff}-A_m(H_{eff}-l_m)} = \frac{\sigma A_m}{H_{eff}}\left(\frac{l_m}{H_{eff}-\frac{A_m}{A_p}(H_{eff}-l_m)}\right) \quad (9)$$

where $G_0$ is the calculated open pore conductance (see Eq. 2) and $H_{eff}$ refers to the effective length of the pore determined from DNA blockage depth, as described in section S1.

By plugging in experimentally measured blockage depths and comparing to the blockages predicted by the resistors-in-series model (as seen in Table S1), we can obtain information about the likely orientation of the translocation molecules. (In order to treat both the DNA and the protein molecule in the same way throughout our calculations, we use the reported crystallographic edge-to-edge distances for both molecules: 2.2 nm for the DNA and 4.2 x 5.6 nm for the MSA). Yusko and Houghtaling and coworkers(4, 5) demonstrated with the electrical shape factor model that the measured blockage depth samples protein orientation, since a non-spherical particle produces a deeper blockage when its longer axis is perpendicular to the electric field in the pore. (6) Further, Ledden and coworkers(7) used a volume-based model to confirm the pore's ability to distinguish the folding state of a protein by comparing protein translocation in native and denaturing conditions.

To test the reliability of the resistors-in-series model, we compare its predictions with experimentally measured values, alongside the values predicted by the established electrical shape factor model for a well-known structure: fully folded tetrameric streptavidin. (As a sanity check, the resistor-in-series model was first applied to the analysis of 1000 bp DNA translocations used to determine pore length. If we section the DNA strand into cylindrical segments of arbitrary length stacked end-to-end within the pore, the predicted blockage depth values of the resistor-in-series model correspond to the experimentally observed blockage depths, offering preliminary validation of the resistors-in-series model.) In Table S1, a comparison is made between the measured blockage depth and the values obtained for the cylindrical resistors-in-series model, the electrical shape factor model (with a correction factor for the ratio of molecular diameter to pore diameter), and the electrical shape factor model with a spherical particle shape and no correction factor. As seen in the second row of Table S1, the blockages predicted by the electrical shape factor model for tetrameric streptavidin align well with the predicted blockages of the resistors-in-series model. The model predicts values representing the extremes of conductance blockage depth induced by protein orientation, meaning the depths of the majority of events will lie between these extremes, as individual streptavidin molecules present different faces (or sample various orientations(34, 36, 42)) during their passage through the pore. The measured blockage depth amplitude and spread (19 ± 5 nS) correspond well with the values generated by the resistors-in-series model, confirming the reliability of this approach.

Interestingly, the cylindrical resistors-in-series model requires only the crystallographic axes of the protein to generate the maximum and minimum expected blockages, whereas the electrical shape factor model requires a knowledge of the volume, axes and orientation of the protein.

| Measured blockage | Cylindrical resistors-in-series model $\Delta G = \frac{\sigma A_m}{H_{eff}}\left(\frac{l_m}{H_{eff} - \frac{A_m}{A_p}(H_{eff} - l_m)}\right)$ | | | | Electrical shape factor model $\Lambda(t) = \frac{(\Delta I/I_0)A_p H_{eff}}{\gamma}\left(1 - 0.8\left(\frac{d_m}{d_p}\right)^3\right)$ | | Electrical shape factor model (no $d_m/d_p$ correction factor) $\Lambda(t) = \frac{(\Delta I/I_0)A_p H_{eff}}{(\gamma = 1.5)}$ | |
|---|---|---|---|---|---|---|---|---|
| | Expected blockage | Molecular dimensions (diameter x length) | Schematic 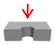 | Proposed molecule structure | Expected blockage | Molecular description | Expected blockage | Molecular description |
| **19 ± 5 nS** | 26 nS | 5.6 nm x 4.2 nm | 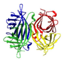 | Tetramer | 20 to 28 nS | Oblate spheroid, γ = 1.42 or 1.70, $d_m$ = 4.2 or 5.6 nm, V = 107 $nm^3$ | 19 nS | Sphere, V = 107 $nm^3$ |
| | 16 nS | 4.2 nm x 5.6 nm | 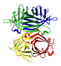 | | 19 to 26 nS | Prolate spheroid, γ = 1.35 or 1.59, $d_m$ = 4.2 or 5.6 nm, V = 107 $nm^3$ | | |
| **7 ± 1 nS** | 9 nS | 4.2 nm x 2.8 nm | 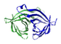 | Dimer(s) | 7 to 9 nS | Oblate spheroid, γ = 1.38 or 1.8, $d_m$ = 4.2 or 2.8 nm, V = 40 $nm^3$ | 7 nS | Sphere, V = 40 $nm^3$ |
| | 7 nS | 2.1 nm x 11 nm | 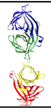 | | 10 nS | Prolate spheroid, γ = 1.05, $d_m$ = 2.1 nm, V = 80 $nm^3$ | | |
| | 5 nS | 2.8 nm x 4.2 nm | 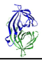 | | 6 to 8 nS | Prolate spheroid, γ = 1.3 or 1.62, $d_m$ = 2.8 or 4.2 nm, V = 40 $nm^3$ | | |
| **3 ± 1 nS** | 4 nS | 2.1 nm x 5.6 nm | 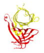 | | 5 to 11 nS | Prolate spheroid, γ = 1.14 or 1.78, $d_m$ = 2.1 or 5.6 nm, V = 40 $nm^3$ | 3 nS | Sphere, V = 18 $nm^3$ |
| | 2 nS | 2.1 nm x 2.8 nm | 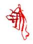 | Monomer | 3 nS | Prolate spheroid, γ = 1.35 or 1.59 $d_m$ = 2.1 or 2.8 nm, V = 18 $nm^3$ | | |

**Table S1 – Table describing expected blockage depths for various dimensions corresponding to streptavidin.**

### S5. Calculation of diffusion coefficient

The He and Niemeyer(8) equation for the diffusion coefficient $D$ is:

$$D = \frac{6.85\times10^{-15}T}{\eta\cdot\sqrt{M^{1/3}r_g}} \tag{10}$$

where $D$ is in $m^2/s$, temperature $T$ is in K, viscosity $\eta$ is in Pa·s, mass $M$ is in kg/kmol, and radius of gyration $r_g$ is in Å. Substituting our values ($T$ = 298 K, $\eta$ of 1350.4 µPa·s for 4 M NaCl,(9) $M$ = 54,500 kg/kmol, $r_g$ = 22 Å(10)) produces a value for the diffusion coefficient of MSA of 52.3 $nm^2/\mu s$ in 4 M NaCl.

For 0.5 M KCl, we perform the same calculation using a reported viscosity value(9) of 898.9 µPa·s to obtain a diffusion coefficient of 78.6 $nm^2/\mu s$.

For 2.9 M LiCl, we have approximated the viscosity by the reported value(11) of 1.29 mPa·s (or in units of $10^{-3}$ $N/m^2$) for 2.6 M LiCl since this is the closest experimental measurement available.(12) The diffusion coefficient is calculated to be 54.8 $nm^2/\mu s$.

#### S6. Calculation of Stokes-Einstein radius

From the diffusion coefficient, we use the Stokes-Einstein relation to find the radius $r_h$ of an equivalent hard sphere diffusing at the same rate as the protein under these experimental conditions (sometimes called the protein's hydrodynamic radius), as follows:

$$r_h = \frac{k_B T}{6\pi\eta D}. \tag{11}$$

Substituting $k_B T$ = 4.114 pN·nm, $\eta$ = 1350.4 x $10^{-6}$ N·s/m$^2$, and $D$ = 5.234 x $10^{-11}$ m$^2$/s, we find the hydrodynamic radius of monovalent streptavidin in 4 M NaCl at 25° C to be $r_h$ = 3.1 nm.

### S7. Calculation of effective charge

Following Muthukumar(13) and Huckel,(14) the effective charge of a spherical particle and its counterion cloud can be described as:

$$Q(r_p) = \frac{Q}{1+\kappa r_g} \tag{12}$$

where $r_g$ is the radius of gyration and $\kappa$ is the inverse Debye length for an aqueous solution at 25°C with a concentration of monovalent salt $C_s$ given in molarity (mol/L) as described by the equation:

$$\kappa = 3.28\sqrt{C_s}\, nm^{-1}. \tag{13}$$

Substituting 4 M for the experimental concentration of NaCl generates a value of 6.56 $nm^{-1}$ for $\kappa$. Multiplying $\kappa$ by $r_g$ (reportedly 2.2 nm for MSA(10)), we obtain a value for $\kappa r_g$ of 14.4. From Eq. 12, the effective charge $Q(r_p)$ is then 6.5% of the original charge value $Q$ or -0.79e.

#### S8. Effect of confinement

Confinement effects reduce protein mobility in pores with diameters less than 1.5 times the protein hydrodynamic diameter, as reported elsewhere.(15–17) In fact, pores smaller than 9.3 nm will likely reduce the mobility of streptavidin with its hydrodynamic radius of 3.1 nm in 4 M NaCl at room temperature (calculated in section S6 above).(18) The effects of hydrodynamic drag alone on the bulk diffusion coefficient *D* for a particle in a cylindrical pore can be obtained from the Bowen, Mohammad and Hilal expression(18, 19) valid for a ratio λ of the hydrodynamic radius of the protein to the pore radius ($r_h/r_p$) up to 0.8:

$$D(r_p) = D \times (1 - 2.3\lambda + 1.154\lambda^2 + 0.224\lambda^3). \quad (14)$$

This correction brings the measured in-pore diffusion coefficient $D(r_p)$ of 4.3 x $10^{-2}$ $nm^2/\mu s$ an order of magnitude closer to streptavidin's bulk diffusion coefficient *D* of 52.3 $nm^2/\mu s$ in 4 M NaCl. In fact, a reduction from the bulk diffusion coefficient by two to three orders of magnitude has previously been observed with high-bandwidth nanopore detection.(17)

#### S9. Comparison of MSA translocation at applied voltage of 200 mV versus 800 mV through the same pore

We present typical traces observed for MSA translocations through a 9.6 nm pore in a $SiN_x$ membrane with an effective thickness of 11 nm. Following CBD-based pore fabrication, a ~10 nM solution of MSA was loaded into the *cis* side of the fluidic cell reservoir. Upon application of 200 mV on the *trans* side, we observed virtually no events in 5 minutes of collected current trace (Figure S4a, red), likely because the MSA molecules pass too rapidly through the pore to allow detection at standard bandwidth.

Application of 800 mV produced events in the current trace (Figure S4a, blue) at a rate of 0.4 Hz/nM with blockage depths of roughly 4 ± 1 nS and dwell times centered around 30 ± 20 µs (Figure S4b). The 4-nS blockage depth corresponds to a volume of roughly a third of the full tetrameric streptavidin, suggesting that the streptavidin is separated into dimers at 800 mV in 2 M KCl.

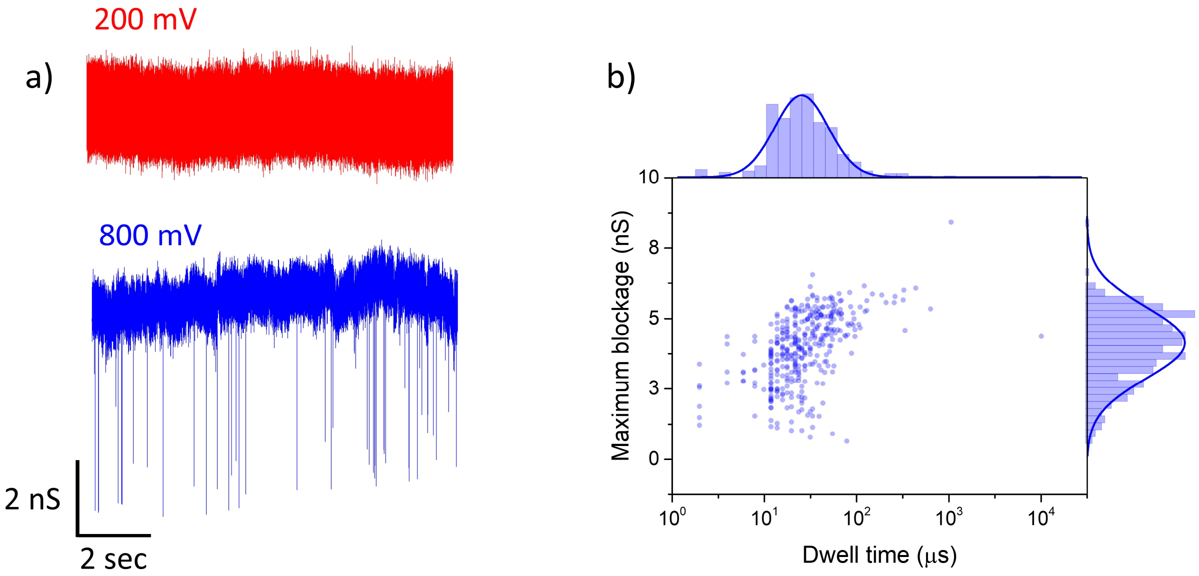


**Figure S4 – Influence of voltage on MSA characteristics in 2 M KCl.** a) Typical 10-second current traces showing nanopore sensing with no events visible at an applied bias of 200 mV (red) for 12 nM MSA and numerous events visible at 800 mV (blue) for 8 nM MSA through a 9.6 nm pore (11 nm thickness). b) Overlay of dwell time versus maximum blockage histograms derived from translocation signals of MSA at 800 mV.

### S10. Calculation of capture radius

Using the equation $r_{eff} = R_C/(2\pi DC)$, where $R_c$ is the detected capture rate, $C$ is the protein concentration, $D$ is the bulk diffusion coefficient, we substitute the experimental values for 4 M NaCl ($R_C$ = 3.13 $s^{-1}$, C= 30 nM or 1.8 $x10^{19}$ molecules/$m^3$, $D$ = 52.3 $x10^{-12}$ $m^2/s$), to arrive at an effective radius of 0.5 nm.

For 0.5 M KCl, we find the effective capture radius of 0.84 nm from $r_{eff} = (300\, molecules \cdot s^{-1})/(2\pi \times 7.86 \times 10^{-1}\ m^2/s \times 7.22 \times 10^{20} molecules/m^3)$.

For 2.9 M LiCl, we find the effective capture radius for the experiment is 0.31 nm from $r_{eff} = (6.4\, molecules \cdot s^{-1})/(2\pi \times 5.479 \times 10^{-11} \mathrm{m^2/s}\ \times 5.9 \times 10^{19} molecules/m^3)$.

### S11. Discrete intra-event levels in KCl and LiCl at 200 mV

For a 7.7 nm pore of 8.7 nm thickness in 2.9 M LiCl (~14.9 S/m), 30 nM MSA-DNA produces a mean maximum blockage of 36 ± 2 nS with 8000 µs mean dwell time at 200 mV (Figure S5). Long intra-event levels occur near 17 nS and 29 nS.

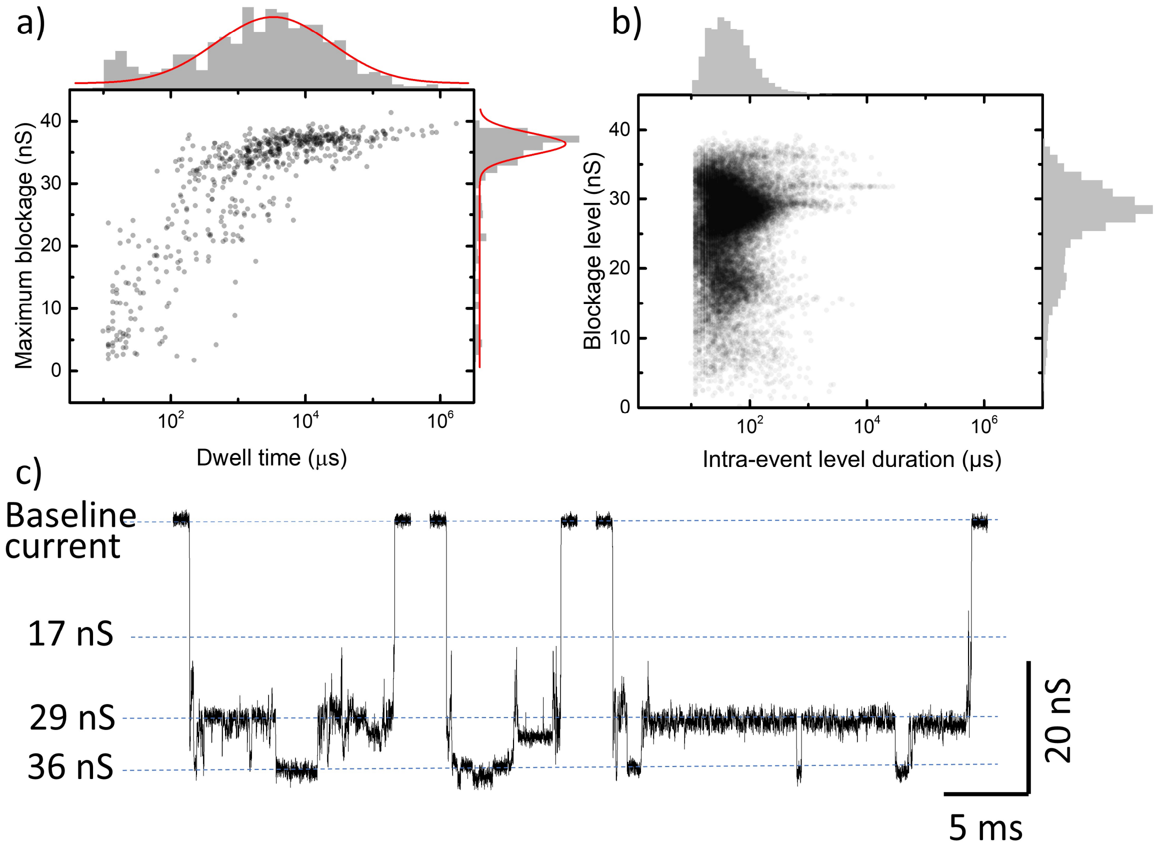


**Figure S5 – Nanopore detection and analysis of MSA bound to biotinylated DNA in 2.9 M LiCl through a 7.7 nm pore (8.7 nm thickness) at 200 mV.** a) Overlay of dwell time and maximum blockage histograms. b) Overlay of intra-event level duration and blockage level histograms for the same events. c) Sample event traces for the same data.

For an 8.5 nm pore with 10.3 nm thickness in 2 M KCl, 8 nM MSA-DNA produced a mean maximum blockage depth of 31 ± 2 nS with a mean dwell time peak near 970 ms (Figure S6). Long intra-event levels occur near 17 nS, with peaks reaching around 30 nS.

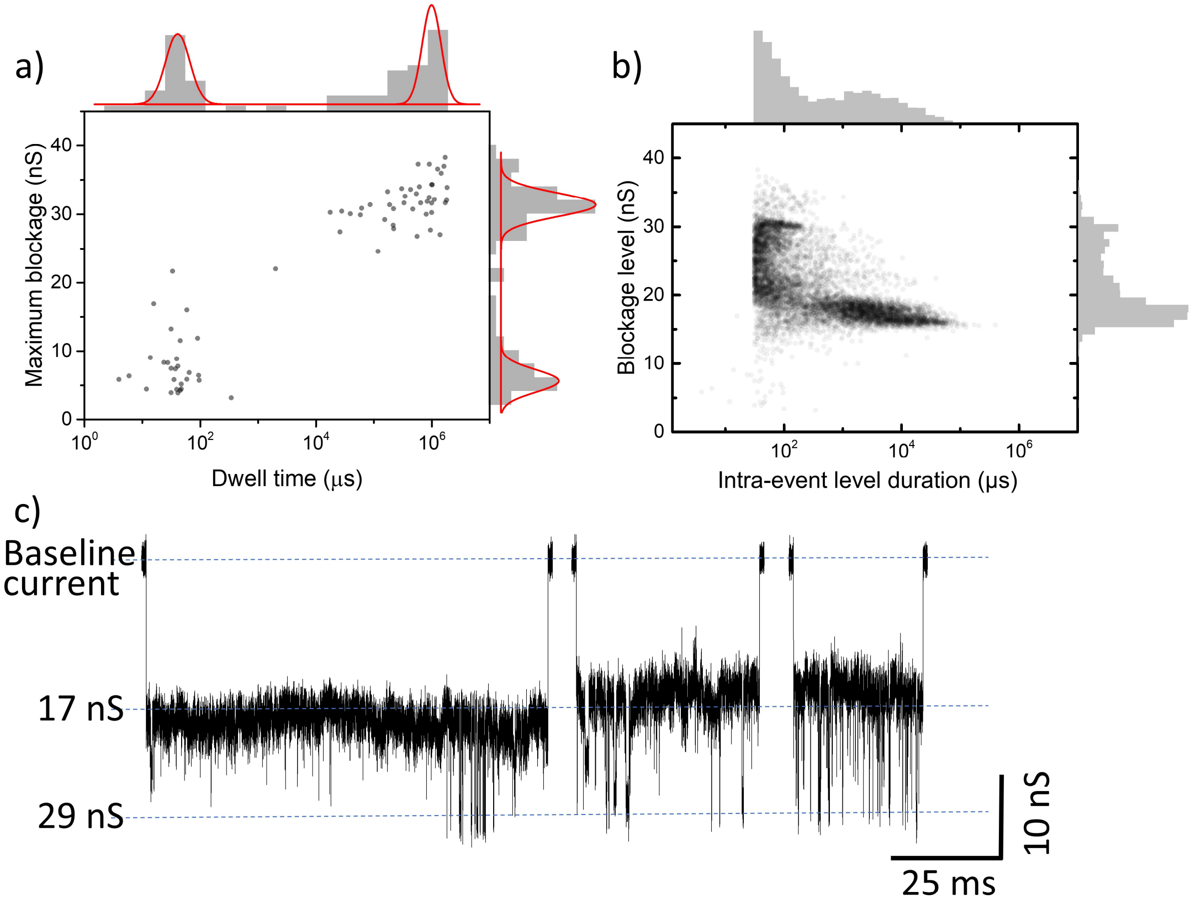


**Figure S6 – Nanopore detection and analysis of MSA bound to biotinylated DNA in 2 M KCl through an 8.5 nm pore (10.3 nm thickness) at 200 mV.** a) Overlay of dwell time and maximum blockage histograms. b) Overlay of intra-event level duration versus blockage level for the same events. c) Sample event traces for the same data.

### S12. Calculation of resistors-in-series model for two macromolecules

#### Case 1: Two macromolecules in series within the pore

In the case where two distinct molecules are present, the resistance of the pore with the molecule inside ($R_{pore\ with\ molecules\ 1\ and\ 2}$) can be expressed as:

$$R_{pore\ with\ molecules\ 1\ and\ 2} = R_{empty\ portion} + R_{portion\ with\ 1} + R_{portion\ with\ 2}$$

$$= \frac{1}{\sigma}\frac{(H_{eff}-l_1-l_2)}{A_p} + \frac{1}{\sigma}\frac{l_1}{A_p-A_1} + \frac{1}{\sigma}\frac{l_2}{A_p-A_2} = \frac{1}{\sigma}\frac{(H_{eff}-l_1-l_2)(A_p-A_2)(A_p-A_1)+A_pl_1(A_p-A_{m2})+A_pl_2(A_p-A_1)}{A_p(A_p-A_2)(A_p-A_1)} \quad (15)$$

where $l_1$ is the length of molecule 1, $l_2$ is the length of molecule 2, $A_1$ is the cross-sectional area of molecule 1, and $A_2$ is the cross sectional area of molecule 2. Inverting the resistance to obtain the conductance, we obtain:

$$R^{-1}_{pore\ with\ molecules\ 1\ and\ 2} = G_{pore\ with\ molecules\ 1\ and\ 2}$$

$$= \frac{\sigma A_p(A_p-A_2)(A_p-A_1)}{(H_{eff}-l_1-l_2)(A_p-A_2)(A_p-A_1)+A_pl_1(A_p-A_2)+A_pl_2(A_p-A_1)}. \quad (16)$$

The conductance change is equal to $\Delta G = G_0 - G_{pore\ with\ molecules\ 1\ and\ 2}$ and gives:

$$\Delta G = \frac{\sigma A_p}{H_{eff}} - \frac{\sigma A_p(A_p-A_2)(A_p-A_1)}{(H_{eff}-l_1-l_2)(A_p-A_2)(A_p-A_1)+A_pl_1(A_p-A_2)+A_pl_2(A_p-A_1)}. \quad (17)$$

An example of this type of calculation can be seen in Table S2 where an expected blockage of 32 nS corresponds to DNA trailing behind streptavidin.

| Measured maximum blockage | Cylindrical resistors-in-series model (molecule → pore) | | | |
|---|---|---|---|---|
| | Expected blockage | Molecular dimensions | | Schematic of molecule |
| | | $R_1$ ($d_1$ x $l_1$) | $R_2$ ($d_2$ x $l_2$) | $R_1$ $R_2$ |
| 52 ± 1 nS | 53 nS | 6.6 x 6.5 nm | --- | 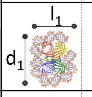 |
| | 50 nS | 6.6 x 5.5 nm | 2.2 x 5.5 nm | 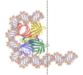 |
| 45 ± 5 nS | 42 nS | 6.2 x 5.5 nm | 2.2 x 5.5 nm | 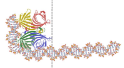 |
| 37 ± 4 nS | 38 nS | 6.2 x 4.5 nm | 2.2 x 6.5 nm | 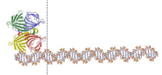 |
| | 32 nS | 5.8 x 4.5 nm | 2.2 x 6.5 nm | 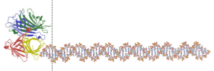 |
| 27 ± 1 nS | 27 nS | 5.0 x 5.8 nm | 2.2 x 5.2 nm | 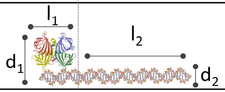 |

**Table S2 – Table comparing measured blockage depths and expected blockage depths derived from cited molecular dimensions using the cylindrical resistors-in-series model.**

##### Case 2: Two overlapping macromolecules (e.g. side-by-side) within the pore

In the case where molecules enter the pore side by side, such as MSA alongside DNA, we can express the combined cross-sectional area of the two molecules then extract an effective radius as follows:

$$A_{MSA} + A_{DNA} = \pi r_{MSA}^2 + \pi r_{DNA}^2 = \pi(r_{MSA}^2 + r_{DNA}^2) = \pi r_{eff}^2 \therefore r_{eff} = \sqrt{r_{MSA}^2 + r_{DNA}^2}. \quad (18)$$

For the DNA alongside widthwise streptavidin, we obtain a diameter of 6.2 nm and a height of 4.5 nm. To test the reasonableness of the resistors-in-series approach, we compare the value it generates to that produced by the electrical shape factor model for a molecule with the same dimensions. The resistors-in-series approach gives 38 nS for this arrangement, while the electrical shape factor approach gives 36 nS (6.2 nm diameter, $\gamma$= 1.5, 150 $nm^3$ volume for 133 $nm^3$ MSA and 4.5 nm of DNA for 17.1 $nm^3$ volume). Examples of this type of calculation can be seen in Table S2 for expected blockages of 27 and 38 nS.

(For the DNA alongside lengthwise streptavidin, we obtain a diameter of 5.0 and a height of 5.8 nm. The cylindrical resistors in series approach gives 27 nS, while the electrical shape factor approach gives 29 nS [5 nm diameter, $\gamma_{para}$ of 1.42 for prolate with diameter 5 nm and length 5.8 nm, 155 $nm^3$ volume for 5.8 nm of DNA + 133 $nm^3$ of MSA]).

##### Case 3: One macromolecule bending around another macromolecule

When the DNA bends around the streptavidin, finding the length of the complex is less straightforward. We explore this possibility using three approaches. The rectangular-prism/cylindrical resistors-in-series method (1) and electrical shape factor method (2) are used to verify the reasonableness of the cylindrical resistor-in-series method (3).

(1) First, we apply the rectangular-prism/cylindrical resistors-in-series model, treating the DNA as a simple rectangular prism when its length is perpendicular to the electrical field, while treating it as a cylinder when its length is parallel to the field, as seen in Table S3. When alongside the MSA, the DNA contributes to the diameter of the MSA according to Eq. 18.

When the MSA advances in the pore and causes the tail end of the DNA to curl around it (see Table S3, 42 nS expected blockage), we consider the far end of the bent DNA tail (which is perpendicular to the electric field) to be a rectangular prism with length equivalent to the width of the DNA diameter plus the width of the MSA (2.2 nm + 5.8 nm = 8.0 nm) and a height of 2.2 nm. Combining this with the segment of the pore containing the DNA strand alongside the MSA (a cylinder with combined diameter 6.2 nm and height 4.5 nm) and the segment of the pore containing the extended cylindrical DNA strand alone, we find an expected blockage depth of 42 nS (Table S3).

Values obtained for other conformations using this approach are shown in Table S3. We note that the DNA length perpendicular to the electric field is greater than the nanopore diameter (8.6 nm) for the 50 nS and 53 nS case. The dimensions used here approximate the molecule area perpendicular to the electric field, with the goal of comparison to the values generated by the cylindrical resistors-series-model for these complex MSA-DNA conformations.

| Measured blockage | Resistors-in-series model with DNA as rectangular prism (if perpendicular to electric field) or as cylinder | | | | | |
|---|---|---|---|---|---|---|
| | Expected blockage | Molecular dimensions | | | Molecule (side-view) | Schematic 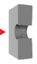  |
| | | R1 ($w \times l$) | R2 ($d \times l$) | R3 | | R1 R2 R3 |
| 52 ± 1 nS | 53 nS | 10.2 x 1.95 | 6.6 x 4.5 | 10.2 w x 1.95 l | 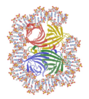 | 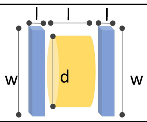  |
| | 50 nS | 10.2 x 1.95 | 6.6 x 4.5 | 2.2 d x 4.55 l | 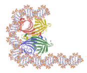 | 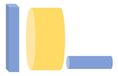 |
| 45 ± 5 nS | 42 nS | 8.0 x 1.95 | 6.2 x 4.5 | 2.2 d x 4.55 l | 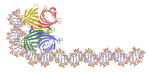 | 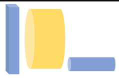 |
| 37 ± 4 nS | 38 nS | 5.5 x 5.8 | 8.9 x 1.95 | 2.2 d x 3.25 l | 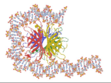 | 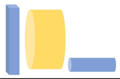 |
| 27 ± 1 nS | 30 nS | 5.0 x 5.8 | 6.7 x 1.95 | 2.2 d x 3.25 l | 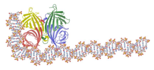 | 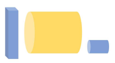 |

**Table S3 – Table compared measured blockage depths with expected blockage depths derived from cited molecular dimensions using the resistors-in-series model with DNA treated as a rectangular prism when situated with its length perpendicular to the electric field or as a cylinder when its length is parallel to the electric field.**

(2) In a second approach, we look at the same conformation (i.e., DNA bending around width-wise streptavidin) and treat the problem using the electrical shape factor method (Table S4). We consider the effective diameter of the double-stranded DNA alongside the streptavidin to be 6.2 nm following Eq. 18 and assume roughly 10.4 nm of the total DNA length is engaged with the streptavidin (calculated by adding the 4.5 nm and 5.8 nm axes of MSA), suggesting a DNA volume of 39 $nm^3$. Adding this to the MSA volume of 133 $nm^3$ gives a total volume of 172 $nm^3$. A spherical particle with these dimensions would give 40.9 nS by the electrical shape factor approach. (A prolate spheroid of the same volume with diameter 6.2 nm and length 6.7 nm [amounting to the 4.5 nm length of the MSA + the 2.2 nm height of the DNA lying perpendicular to the electric field] would give a maximum conductance blockage of 47 nS and a minimum of 40 nS with correction factor included, where $m$ = 1.08, $\gamma_{para}$ = 1.45 and $\gamma_{perp}$ = 1.52).

<table>
<tr><th rowspan="2">Measured blockage</th><th colspan="2">Electrical shape factor model<br>$\Lambda(t) = \frac{(\Delta I/I_0)A_p H_{eff}}{\gamma}\left(1 - 0.8\left(\frac{d_m}{d_p}\right)^3\right)$</th><th colspan="3">Electrical shape factor model<br>(no $d_m/d_p$ correction factor)<br>$\Lambda(t) = \frac{(\Delta I/I_0)A_p H_{eff}}{(\gamma = 1.5)}$</th></tr>
<tr><th>Expected blockage</th><th>Molecular description</th><th>Expected blockage</th><th>Molecular description</th><th>Volume explanation</th></tr>
<tr><td>52 ± 1 nS</td><td>52 to 54 nS</td><td>Oblate spheroid,<br>γ = 1.49 or 1.51,<br>$d_m$ = 6.6 or 6.5 nm,<br>V = 205 $nm^3$</td><td>53 nS</td><td>Sphere,<br>$d_m$= 6.6 nm,<br>V = 205 $nm^3$</td><td>133 $nm^3$ (MSA) + 72 $nm^3$<br>(full DNA length of 19 nm)</td></tr>
<tr><td rowspan="2">45 ± 5 nS</td><td rowspan="3">40 to 47 nS</td><td rowspan="3">Prolate spheroid,<br>γ = 1.45 or 1.52,<br>$d_m$ = 6.2 or 6.7 nm,<br>V = 172 $nm^3$</td><td>49 nS</td><td>Sphere,<br>$d_m$= 6.6 nm,<br>V = 189 $nm^3$</td><td>133 $nm^3$ (MSA)<br>+ 56 $nm^3$ (14.8 nm length of DNA)</td></tr>
<tr><td>41 nS</td><td>Sphere,<br>$d_m$= 6.2 nm<br>V = 172 $nm^3$</td><td>133 $nm^3$ MSA<br>+ 39 $nm^3$ (10.3 nm length of DNA)</td></tr>
<tr><td>37 ± 4 nS</td><td>36 nS</td><td>Sphere,<br>$d_m$= 6.2 nm,<br>V= 150 $nm^3$</td><td>133 $nm^3$ MSA<br>+ 17.1 $nm^3$<br>(4.5 nm length of DNA)</td></tr>
<tr><td>27 ± 1 nS</td><td>29 nS</td><td>Prolate spheroid,<br>γ = 1.42,<br>$d_m$ = 5 nm, V = 155 $nm^3$</td><td>31 nS</td><td>Sphere,<br>$d_m$= 5.0 nm,<br>V = 155 $nm^3$</td><td>133 $nm^3$ MSA<br>+ 22 $nm^3$<br>(5.8 nm length of DNA)</td></tr>
</table>

**Table S4 – Table comparing measured blockage depths with expected blockage depths for various spheroids using the electrical shape factor model.**

(3) Finally, we apply the cylindrical resistors-in-series Eq. 17 to the same conformation – DNA bending around widthwise streptavidin – and section the pore according to the various portions of the complex. Now, $A_1$ corresponds to the cross-sectional area of the DNA curled over the MSA while $l_1$ describes the length of the complex that maintains that same cross-sectional area. $A_2$ and $l_2$ describe the remaining cross-sectional area and length of the DNA tail. (As before, $A_p$ and $H_{eff}$ describe the area and the DNA-calibrated length of the pore, respectively.)

$$\Delta G = \frac{\sigma A_p}{H_{eff}} - \frac{\sigma A_p(A_p - A_2)(A_p - A_1)}{(H_{eff} - l_1 - l_2)(A_p - A_2)(A_p - A_1) + A_p l_1(A_p - A_2) + A_p l_2(A_p - A_1)} \tag{19}$$

To obtain a similar blockage depth to the values generated by the rectangular-prism/cylindrical resistors-in-series model (Table S3) and the electrical shape factor model described above, we consider the total length of the complex with the DNA roughly halfway around the streptavidin as 5.5 nm, as this produces an expected blockage depth of 42 nS (see Table S2).

Similarly, when the DNA curls further around the MSA, the rectangular-prism/cylindrical resistors-in-series approach gives 48 nS (10.2 nm prism atop a 6.6 nm diameter molecule with 4.5 nm height), while the electrical shape factor approach also gives 49 nS (6.6 nm diameter molecule, 186 $nm^3$ volume for 133 $nm^3$ of MSA plus 14.8 nm of DNA, $\gamma = 1.5$). This corresponds well with the 48 nS value found using the simple cylindrical resistors-in-series approach (6.6 nm diameter, 5.5 nm length).

If the DNA wraps completely around the MSA, the electrical shape factor approach generates a value of 53 nS (6.6 nm sphere, 205 $nm^3$ volume for MSA-DNA). The same value is generated by the resistors-in-series method treating the top and bottom sections of the DNA strand as rectangular prisms (two 2.2 nm x 10.2 nm prisms for DNA surrounding a 6.6 nm x 4.5 nm cylinder). For this reason, we select a length of 6.5 nm in the cylindrical resistors-in-series model (6.6 nm diameter, 6.5 nm length) to produce a corresponding blockage depth of 53 nS.

#### REFERENCES


1. Kowalczyk, S.W., A.Y. Grosberg, Y. Rabin, and C. Dekker. 2011. Modeling the conductance and DNA blockade of solid-state nanopores. *Nanotechnology*. 22:315101.

2. Grover, N.B., J. Naaman, S. Ben-Sasson, and F. Doljanski. 1969. Electrical Sizing of Particles in Suspensions: I. Theory. *Biophys. J.* 9:1398–1414.

3. Golibersuch, D.C. 1973. Observation of Aspherical Particle Rotation in Poiseuille Flow via the Resistance Pulse Technique: I. Application to Human Erythrocytes. *Biophys. J.* 13:265.

4. Yusko, E.C., B.R. Bruhn, O.M. Eggenberger, J. Houghtaling, R.C. Rollings, N.C. Walsh, S. Nandivada, M. Pindrus, A.R. Hall, D. Sept, J. Li, D.S. Kalonia, and M. Mayer. 2017. Real-time shape approximation and fingerprinting of single proteins using a nanopore. *Nat. Nanotechnol.* 12:360–367.

5. Houghtaling, J., C. Ying, O.M. Eggenberger, A. Fennouri, S. Nandivada, M. Acharjee, J. Li, A.R. Hall, and M. Mayer. 2019. Estimation of Shape, Volume, and Dipole Moment of Individual Proteins Freely Transiting a Synthetic Nanopore. *ACS Nano*. 13:5231–5242.

6. Freedman, K.J., S.R. Haq, J.B. Edel, P. Jemth, and M.J. Kim. 2013. Single molecule unfolding and stretching of protein domains inside a solid-state nanopore by electric field. *Sci. Rep.* 3:1638.

7. Li, J., D. Fologea, R. Rollings, and B. Ledden. 2014. Characterization of Protein Unfolding with Solid-state Nanopores. *Protein Pept. Lett.* 21:256–265.

8. He, L.-Z., and B. Niemeyer. 2003. A Novel Correlation for Protein Diffusion Coefficients Based on Molecular Weight and Radius of Gyration. *Biotechnol. Prog.* 19:544–548.

9. Kestin, J., H.E. Khalifa, and R.J. Correia. 1981. Tables of the dynamic and kinematic viscosity of aqueous NaCl solutions in the temperature range 20-150 °C and the pressure range 0.1-35 MPa. *J. Phys. Chem. Ref. Data*. 10:71.

10. Haridasan, N., S.K. Kannam, S. Mogurampelly, and S.P. Sathian. 2018. Translational mobilities of proteins in nanochannels: A coarse-grained molecular dynamics study. *Phys. Rev. E*. 97:062415.

11. Abdulagatov, I.M., A.B. Zeinalova, and N.D. Azizov. 2006. Experimental viscosity B-coefficients of aqueous LiCl solutions. *J. Mol. Liq.* 126:75–88.

12. Mao, S., Z. Duan, S. Mao, and Z. Duan. 2009. The Viscosity of Aqueous Alkali-Chloride Solutions up to 623 K, 1,000 bar, and High Ionic Strength. *Int J Thermophys*. 30:1510–1523.

13. Muthukumar, M. 2014. Communication: Charge, diffusion, and mobility of proteins through nanopores. *J. Chem. Phys.* 141:081104.

14. E. Huckel. 1924. Die kataphorese der kugel. *Phys.Z.* 25:204–210.

15. Talaga, D.S., and J. Li. 2009. Single-molecule protein unfolding in solid state nanopores. *J. Am. Chem. Soc.* 131:9287–97.

16. Plesa, C., S.W. Kowalczyk, R. Zinsmeester, A.Y. Grosberg, Y. Rabin, and C. Dekker. 2013. Fast translocation of proteins through solid state nanopores. *Nano Lett.* 13:658–663.

17. Larkin, J., R.Y. Henley, M. Muthukumar, J.K. Rosenstein, and M. Wanunu. 2014. High-

Bandwidth Protein Analysis Using Solid-State Nanopores. *Biophys. J.* 106:696–704.

18. Kannam, S.K., and M.T. Downton. 2017. Translational diffusion of proteins in nanochannels. *J. Chem. Phys.* 146:054108.

19. Bowen, W.R., A.W. Mohammad, and N. Hilal. 1997. Characterisation of nanofiltration membranes for predictive purposes — use of salts, uncharged solutes and atomic force microscopy. *J. Memb. Sci.* 126:91–105.
